# GroEL/S helps purge deleterious mutations and reduce genetic diversity during adaptive protein evolution

**DOI:** 10.1101/2021.03.05.434078

**Authors:** Bharat Ravi Iyengar, Andreas Wagner

## Abstract

Chaperones are proteins that help other proteins fold. They also affect the adaptive evolution of their client proteins by buffering deleterious mutations and increasing the genetic diversity of evolving proteins. We study how the bacterial chaperone GroE (GroEL + GroES) affects the evolution of green fluorescent protein (GFP). To this end we subjected GFP to multiple rounds of mutation and selection for its color phenotype in four replicate *E. coli* populations, and studied its evolutionary dynamics through high-throughput sequencing and mutant engineering. We evolved GFP both under stabilizing selection for its ancestral (green) phenotype, and to directional selection for a new (cyan) phenotype,. We did so both under low and high expression of the chaperone GroE. In contrast to prevailing wisdom, we observe that GroE does not just buffer but also helps purge deleterious mutations from evolving populations. In doing so, GroE helps reduce the genetic diversity of evolving populations. In addition, it causes phenotypic heterogeneity in mutants with the same genotype, potentiating their effect in some cells, and buffering it in others. Our observations show that chaperones can affect adaptive evolution through more than one mechanism.

**Highlights:**

- GroE reduces genetic diversity
- GroE potentiates the effect of deleterious mutations
- GroE intensifies purifying selection and leads to higher activity of client proteins

## Introduction

In most proteins, the majority of amino acids help provide a stable structural scaffold, whereas fewer amino acids are directly responsible for catalysis or other protein activities^1^. Protein evolution is thus constrained by mutations that destabilize a protein’s three-dimensional fold^2,3^. Such mutations can reduce protein activity and organismal fitness, for example by reducing the amount of correctly folded and thus active protein. They can also increase a protein’s propensity to form toxic aggregates of misfolded proteins^4–6^. In most proteins, the majority of amino acids help provide a stable structural scaffold, whereas fewer amino acids are directly responsible for catalysis or other protein activities^1^. Mutations that create a new protein activity are especially often destabilizing^7–10^.

Cells encode multiple proteins called chaperones that are dedicated to help other proteins to fold correctly and to maintain their fold. Chaperones act via various mechanisms, such as the stabilization of newly synthesized polypeptides, the acceleration of the folding process, and the refolding of misfolded proteins. This diversity of mechanisms is reflected in a diversity of chaperone structures^11–13^. Prominent chaperone classes include the protein family Hsp60 (heat shock protein with a molecular weight of 60Kda), the Hsp70, Hsp90 and Hsp100 families, as well as the trigger factor. Chaperones from all these families exist in both bacteria and eukaryotes^11–13^.

The GroEL/S complex (GroE) is one of the major chaperones in bacteria. It is composed of the essential proteins GroEL and GroES^14,15^, and belongs to the Hsp60 family. Eukaryotes also express a GroE homolog, which helps mitochondrial and chloroplast proteins fold. Structurally, GroE belongs to a class of chaperones known as chaperonins, which form a cylindrical cage that entraps an unfolded polypeptide molecule and allows it to refold^16^.

During adaptive evolution, chaperones can facilitate the evolution of various organismal traits, including the evolution of proteins with new functions^17–23^. For example, Hsp90 accelerates the evolution of drug resistance in fungi^20^. In addition, chaperones can prevent the erosion of organismal fitness when deleterious mutations accumulate in an evolving population. For example, overexpressing GroE in *E. coli^24^* and *Salmonellla typhimurium^25^* populations with large number of random genomic DNA mutations can improve bacterial population growth. Relatedly, overexpressing GroE in *E.coli* populations subject to periodic bottlenecking reduces the likelihood of population extinction^26^.

A main mechanism by which chaperones may facilitate adaptive evolution is the buffering of deleterious mutations^18,23,24,27^. By helping a protein with a destabilizing mutation fold correctly, a chaperone can buffer the deleterious effects of this mutation. This mechanism is especially well documented for GroE^17,28,29^. For example, GroE directly improves the folding rate and the fluorescence of a green fluorescent protein (GFP) variant whose fluorescence is compromised by the mutation K45E (a lysine [K] to glutamate [E] change at position 45)^29^. Additionally, GroE overexpression can promote the evolution of new protein functions by stabilizing proteins^17,18^. For example, an F306L mutation that improves the catalytic activity of the enzyme phosphotriesterase on a novel substrate destabilizes the protein, but this destabilizing effect can be mitigated by GroE^18^.

Despite the plausibility of this buffering mechanism, several reports on Hsp90 suggest that this chaperone can also have the opposite effect. That is, it can *potentiate* or enhance the effect of a mutation^20,30–32^. For example the oncogenic activity of the viral oncogene v-Src is amplified by Hsp90^30^. More generally, Hsp90 has been reported to both buffer^19,27,33^ and potentiate^20,31,32^ mutational effects. Existing work aiming to distinguish Hsp90-mediated buffering from potentiation focuses on complex morphological traits in the yeast *Saccharomyces cerevisiae^32^.* Here we take a complementary approach by studying the influence of a chaperone on the directed evolution of a single protein. In our system, one can study chaperone effects in greater molecular detail by combining high-throughput sequencing of evolving populations with protein engineering. In addition, we focus on the bacterial chaperone GroE, for which buffering but not potentiation has been demonstrated. Most existing experiments on GroE buffering in individual proteins rely on single protein mutations^28,29^ or small populations of variants^17,18^. In contrast, we maintained large populations of more than 10^5^ evolving proteins in which many variants segregate during multiple rounds of directed evolution.

Specifically, we studied the influence of GroE on the adaptive evolution of green fluorescent protein (GFP) in *E. coli* cells that overexpress GroE. We subjected GFP to directed evolution experiments in which we alternated cycles of (PCR-mediated) mutation with selection imposed by fluorescence-activated cell sorting (FACS), both with and without overexpression of GroE. In phase 1 of our experiments, we performed five rounds (“generations”) of evolution under stabilizing selection on the ancestral green fluorescent phenotype. We followed this phase 1 by a phase 2, in which we imposed directional selection on the new color phenotype of cyan fluorescence during an additional five rounds. We studied both stabilizing and directional selection, because a chaperone might have different effects under different types of selection.

We chose GFP in this study for several reasons. First, its light-emission phenotype can be easily measured at single cell resolution in a high throughput manner using flow cytometry. Second, it allows us to exert selection in a highly controlled manner via FACS. Third, GFP is non-native to the *E.coli* host, and interferes less with the host’s cell physiology, growth, and metabolism than native proteins would. Fourth, GFP is a known GroE client, that is, the chaperone can promote GFP folding^34^.

We studied the genotypic and phenotypic evolution of GFP via high-throughput single molecule real time (SMRT) sequencing, protein engineering, and phenotypic analysis. We focused on a key prediction that distinguishes the buffering and potentiation hypotheses: If a chaperone buffers the deleterious effects of mutations, then it should help increase genetic diversity in a population over time, because some mutations that would otherwise be deleterious would be tolerated in its presence. Conversely, if a chaperone potentiates the effect of such mutations, it should lead to a loss of genetic diversity, because it renders such mutations more deleterious. We note that a chaperone may buffer some mutations and potentiate others. Our experiments show that both buffering and potentiation can occur in the same population, but that potentiation far outweighs buffering in its effects on genetic diversity.

## Results

### Experimental design

To evolve GFP under conditions of varying GroE (GroEL + GroES) expression, we first constructed an *E.coli* plasmid (***Figure S2***) that expresses GFP constitutively, and that allowed us to vary chaperone expression via an arabinose-inducible promoter. With this expression system, we studied GFP evolution at different chaperone expression levels. We note that GroEL and GroES are essential proteins, such that the chromosomal genes *groS* and *groL,* cannot be deleted. Thus, when we refer to GroE expression throughout, we strictly refer to overexpression of GroE from the expression plasmid. Consistent with a previous demonstration that GFP is a client of GroE^34^, we found that chaperone expression affects the fluorescence of our ancestral GFP protein (***Figure S4***).

We performed directed evolution in four replicate populations that overexpressed GroE (condition G^+^) and in four other populations that did not (G^-^). In each round (generation) of evolution and for each population, we introduced random mutations into GFP via error-prone PCR at a rate of ~1 nucleotide substitution per GFP-coding gene, corresponding to approximately 0.95 amino acid changes per GFP protein (Materials and Methods). The size of the population bottleneck during our experiment was ~ 10^5^ individuals, such that genetic drift plays a negligible role on the time scale of the experiment.

We selected cells for survival using FACS (***Figure 1***) under two selection regimes that distinguish phase 1 from the later phase 2 of our experiments. In phase 1 we selected cells for survival that showed the native (ancestral) GFP phenotype of green fluorescence. In phase 2, we selected for the new phenotype of cyan fluorescence. Each phase consisted of five rounds (“generations”) of mutagenesis and selection. In both phases we applied weak rather than strong selection for high fluorescence, because we reasoned that strong selection may favor mutants that fold well on their own, may thus not require chaperone assistance, and would thus subvert the intent of our study. Specifically, for each selection step we only required that cells fluoresce more intensely than the autofluorescence of cells not expressing GFP. Each phase consisted of five rounds (“generations”) of mutagenesis and selection. After each round we recorded the phenotype of surviving cells using flow cytometry, and sequenced population samples using SMRT) sequencing^35^.

**Figure 1.**
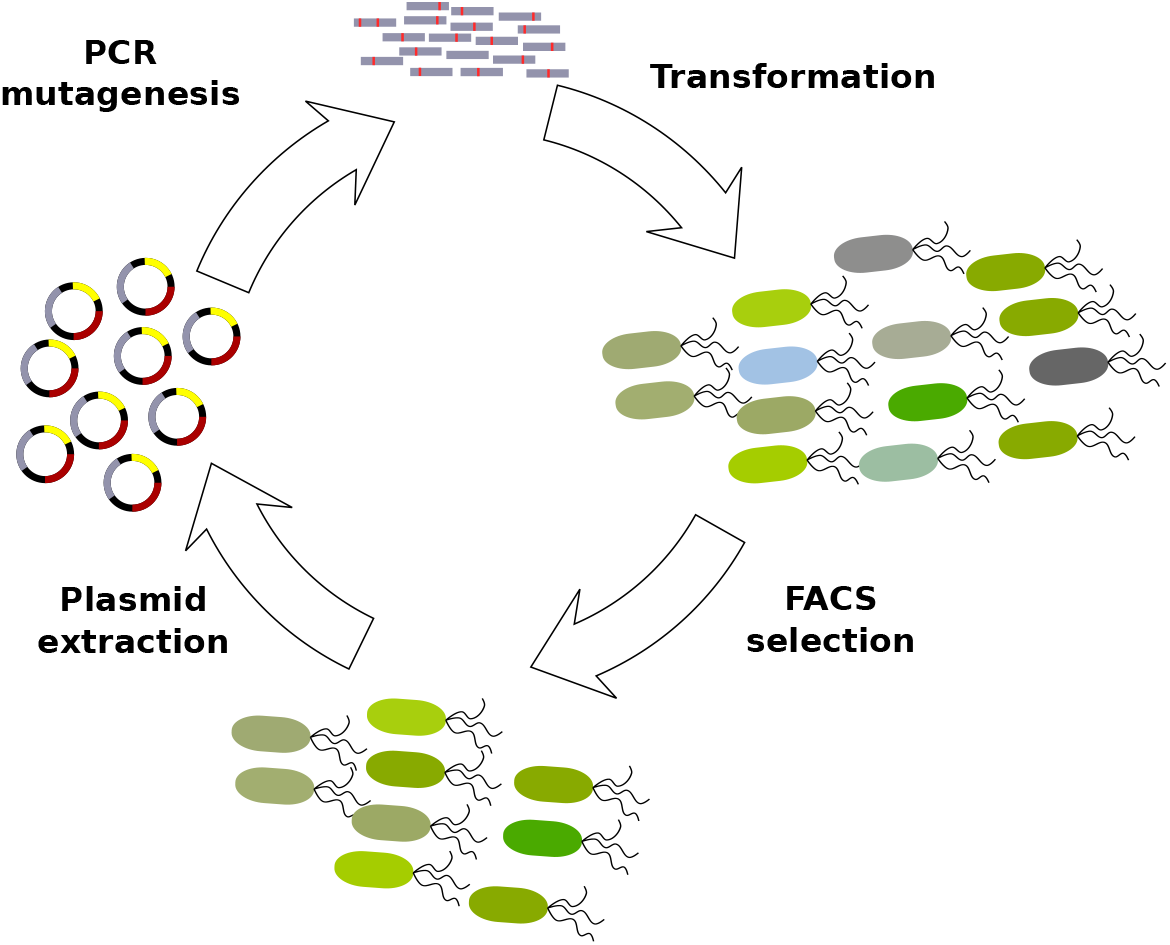
Experimental design. We selected cells for green fluorescence in phase 1 and cyan fluorescence in phase 2. Each phase consisted of five rounds (generations) of directed evolution. We sequenced the GFP gene from plasmids that survived each round of directed evolution using SMRT sequencing.

### GroE expression slows the decay of fluorescence under weak stabilizing selection

The vast majority of mutations affecting protein evolution are deleterious^36–38^. Because phase 1 evolution involved only weak selection on our ancestral green fluorescence phenotype, we would expect that such mutations accumulate in our phase 1 populations. This was indeed the case. We measured the distribution of green fluorescence of 10^5^ single cells from G^+^ and G^-^ populations at the end of each round of phase 1 evolution. During all five generations, green fluorescence consistently declined in all populations relative to the ancestor (***Figure 2***). However, median fluorescence of G^+^ populations declined significantly more slowly than in G^-^ populations *(P =* 10^-7^, linear mixed effects model [LMM], type-III analysis of variance [ANOVA] using Satterthwaite’s method; Materials and Methods). As a result, at the end of phase 1 evolution, all G^+^ populations showed significantly higher median green fluorescence than G^-^ populations (*P* = 0.0088, one tailed Mann-Whitney U-test).

**Figure 2.**
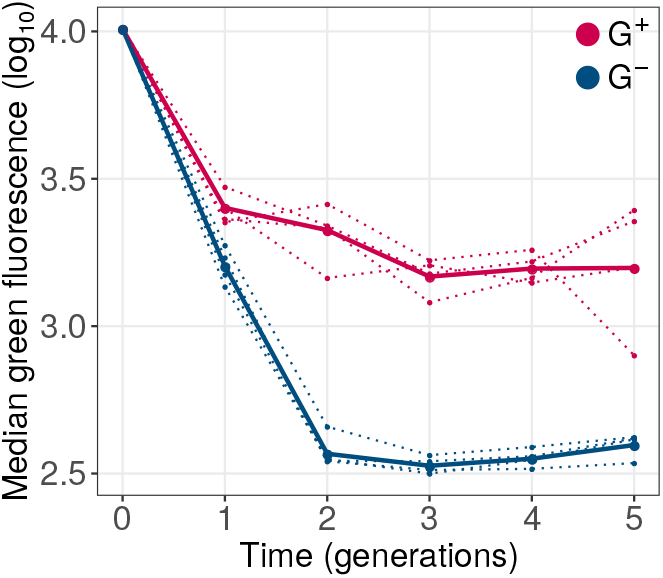
GroE expression reduces the decay of green fluorescence during phase 1 evolution. The vertical axis denotes logarithmically (base 10) transformed median green fluorescence (arbitrary units) of G^+^ (red) and G^-^ (blue) populations. The horizontal axis indicates time in generations (rounds of evolution), with zero referring to the ancestral GFP protein. Dotted lines denote the median fluorescence of individual replicate populations. The solid lines denote the median fluorescence when all four replicate populations are pooled.

### GroE slows genetic diversification under weak stabilizing selection

We next turned to the mechanism by which the chaperone slows down the decay of green fluorescence. If this mechanism relies on buffering the effects of deleterious mutations, then chaperone expression should help increase genetic diversity over time, because some mutations that would otherwise be eliminated by purifying selection could remain in the population. Conversely, if the chaperone acts predominantly by potentiating the effect of deleterious mutations, it should help reduce genetic diversity, because more such mutations would be subject to purifying selection. We define a deleterious mutation as one that reduces fluorescence, because in our experiments selection acts on fluorescence. We note that buffering and potentiation may occur simultaneously in the same population, i.e., GroE may buffer the effect of some mutations while it potentiates the effect of others. To find out which process dominates in its effect on genetic diversity, we sequenced the GFP coding regions from each of the phase 1 populations to a coverage of 1000-3300 (average 2155) single molecule reads, depending on the population. From the sequencing reads, we calculated the frequencies of point mutations and multi-mutant genotypes at the amino acid level.

***Figure 3A*** shows how the mean number of amino acid changes in GFP relative to ancestral GFP, evolves over time. Not surprisingly, both G^+^ and G^-^ populations diverged significantly from the ancestor during evolution (linear mixed effects model [LMM]: ANOVA, *P* < 10^-15^). However, the rate of increase of divergence of G^+^ populations was significantly lower than that of G^-^ populations (LMM: ANOVA, *P* < 10^-5^). We performed analogous analyses for the average pairwise distance between the genotypes in the same population (***Figure 3B***), and for the Shannon entropy (***Figure 3C***), an information-theoretic measure of genetic diversity. We found that both these diversity metrics also increase more slowly in G^+^ populations (LMM: ANOVA, *P* < 0.0012).

**Figure 3.**
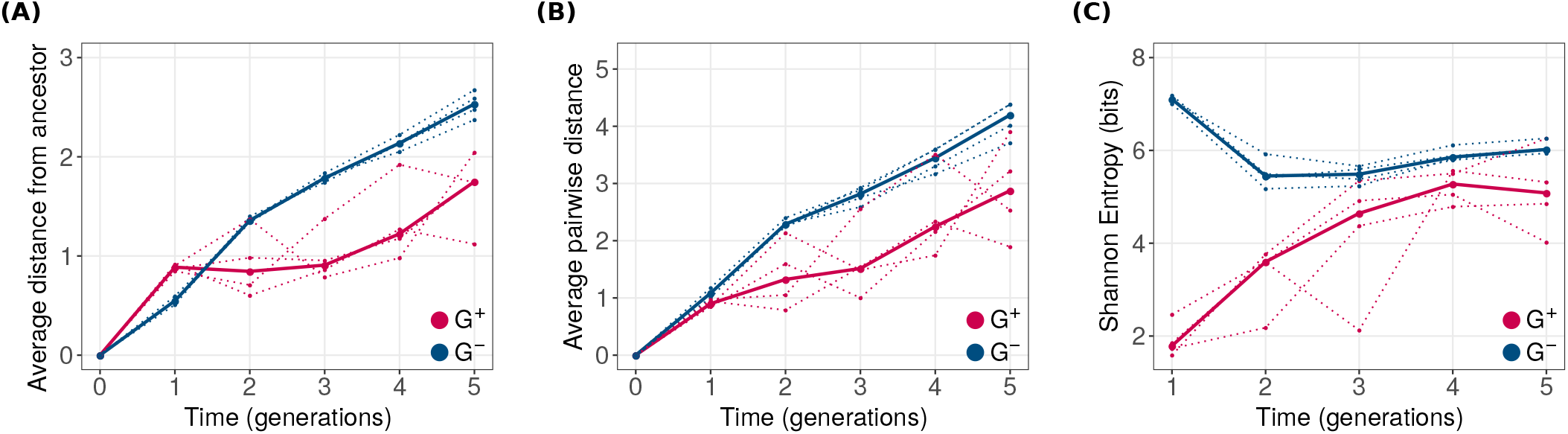
GroE expression leads to reduced genetic diversity during phase 1 evolution. Genetic diversity metrics **(A)** average distance from the ancestral GFP, **(B)** average pairwise distance between genotypes and **(C)** Shannon entropy are shown on the vertical axes. Horizontal axes denote time in generations of evolution, where generation zero corresponds to the ancestral GFP sequence. G^+^ and G^-^ populations are color coded as red and blue, respectively. For all three metrics, G^+^ populations showed significantly lower genetic diversity (LMM: ANOVA, ***P*** < 0.0012).

In sum, GroE reduces genetic diversity in our evolving populations. This supports the view that it predominantly potentiates rather than buffers the effects of deleterious mutations, and thus helps purge such mutations.

In addition to affecting the overall amount of genetic diversity, GroE may cause different kinds of genotypes to accumulate. To find out whether this is the case, we randomly sampled 200 sequences from each population, and displayed the location of these sequences in genotype space using principal component analysis (PCA), a widely used dimensionality reduction method^39^. This analysis shows that G^+^ and G^-^ populations cluster in different regions of genotype space (***Figure S8A***). A complementary principal component analysis on the frequencies of individual amino acid alleles shows analogous differences(***Figure S8B***). Populations evolving with and without GroE expression, harbor different sets of GFP variants.

### GroE buffers the effects of at least some mutations in phase 1 populations

Our preceding analyses do not address the question whether GroE potentiates the effects of all deleterious mutations, or whether it may buffer the effects of at least some mutations. To find out, we focused on another prediction of the buffering hypothesis. If some deleterious mutations that occur in our phase 1 populations are buffered by GroE, these mutations are more likely to remain in the population. Their fluorescence intensity should increase with GroE expression. Thus, if buffering is important, the fluorescence of populations at the end of phase 1 should increase with GroE expression. That is, if we quantify the fluorescence intensity of G^+^ populations during phase 1 in two conditions, one where the chaperone is not overexpressed and one where it is, then fluorescence should be higher when the chaperone is overexpressed. This is not necessarily expected under the potentiation hypothesis, where the chaperone may have simply helped eliminate deleterious mutations, and the remaining mutations may or may not be chaperone dependent. In addition, a potentiating chaperone may also decrease the fluorescence of some of the GFP variants that remain in the final population.

To find out whether fluorescence at the end of phase 1 evolution is chaperone dependent, we measured the fluorescence of those populations that had evolved while GroE was overexpressed, both with and without the induction of the chaperone (***Figure 4***), and compared their median fluorescence using a Mann-Whitney U-test. In three out of four populations chaperone expression increased fluorescence *(P* < 1.5 × 10^-4^), and in one (replicate 3) it decreased fluorescence *(P* < 10^-15^). Although these differences are statistically highly significant because of the large number of individuals we analyzed (*N >* 77000), we also note that they are small in magnitude, ranging from 2 to 11%. They contrast with the much greater differences that emerge in fluorescence during evolution (***Figure 2***), most of which must be caused by potentiation. In sum, some mutational buffering takes place in our evolving populations but its effect on overall fluorescence is small. This conclusion is reinforced by specific candidates for buffered mutants that we engineered and analyzed phenotypically (***SOMsection 8***).

**Figure 4.**
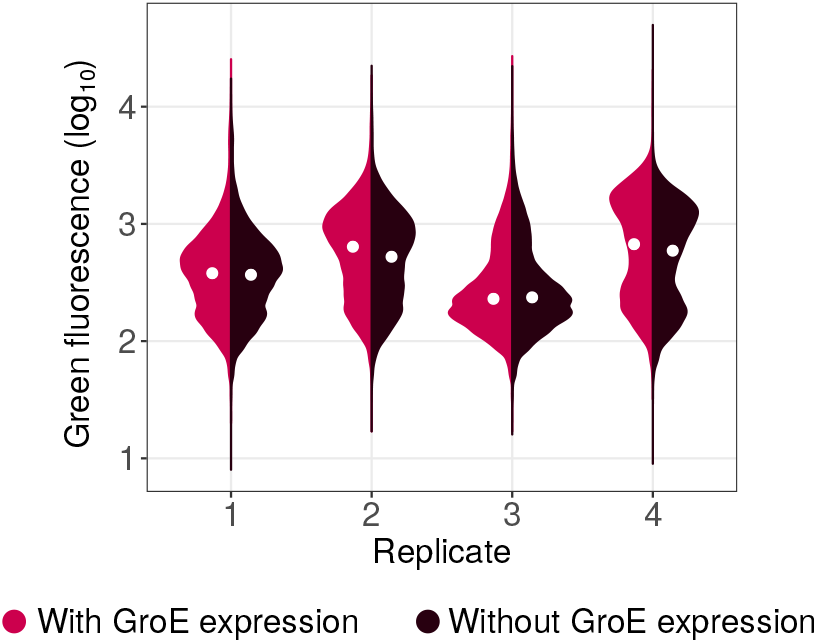
Violin plots denoting the distribution of logarithmically (base 10) transformed green fluorescence (arbitrary units) for each replicate G^+^ population at the end of phase 1 evolution (generation 5), with (red) or without (brown) the expression of GroE. The white circle in the center of the distribution denotes the median. The medians are significantly different for each pair of distributions shown (Mann-Whitney U-test, *P* < 1.5 × 10^−4^).

### GroE disfavors the accumulation of deleterious mutations

To further validate the hypothesis that GroE helps purge deleterious mutations, we examined our sequence data for single amino acid variants that attained significantly lower frequency in G^+^ than in G^-^ populations at the end of phase 1 (Materials and Methods). To keep this analysis tractable, and to restrict ourselves to those mutations that are likely to affect fluorescence most strongly, we restricted this analysis to variants whose frequency exceeded 3.5% at the end of evolution in at least one replicate population (***Figure S10***). We note that this frequency threshold is higher than the expected frequency of any one variant due to mutation pressure alone (*N* = 10^5^, *P* < 10^-5^, Monte-Carlo simulations).

In total we identified seven such variants (GLM:LRT, *P* < 10^-15^ for the null hypothesis that they have equal frequency in G^+^ and G^-^ populations). Specifically, these are the variants M11, M1 L, M1V, S2G, K52R, I128T and N198D. Of these seven variants, the first four had consistently high frequency (8.5 - 67%) in every replicate G^-^ population (***Figure 5A***). More than 87% of individuals in every population had at least one of these four mutations. In contrast, the other three mutations: K52R, I128T and N198D, had comparatively lower frequencies (0.7 - 5.5%; ***Figure S10***). Therefore, we chose to further investigate the mutations M1I, M1L, M1V and S2G.

**Figure 5.**
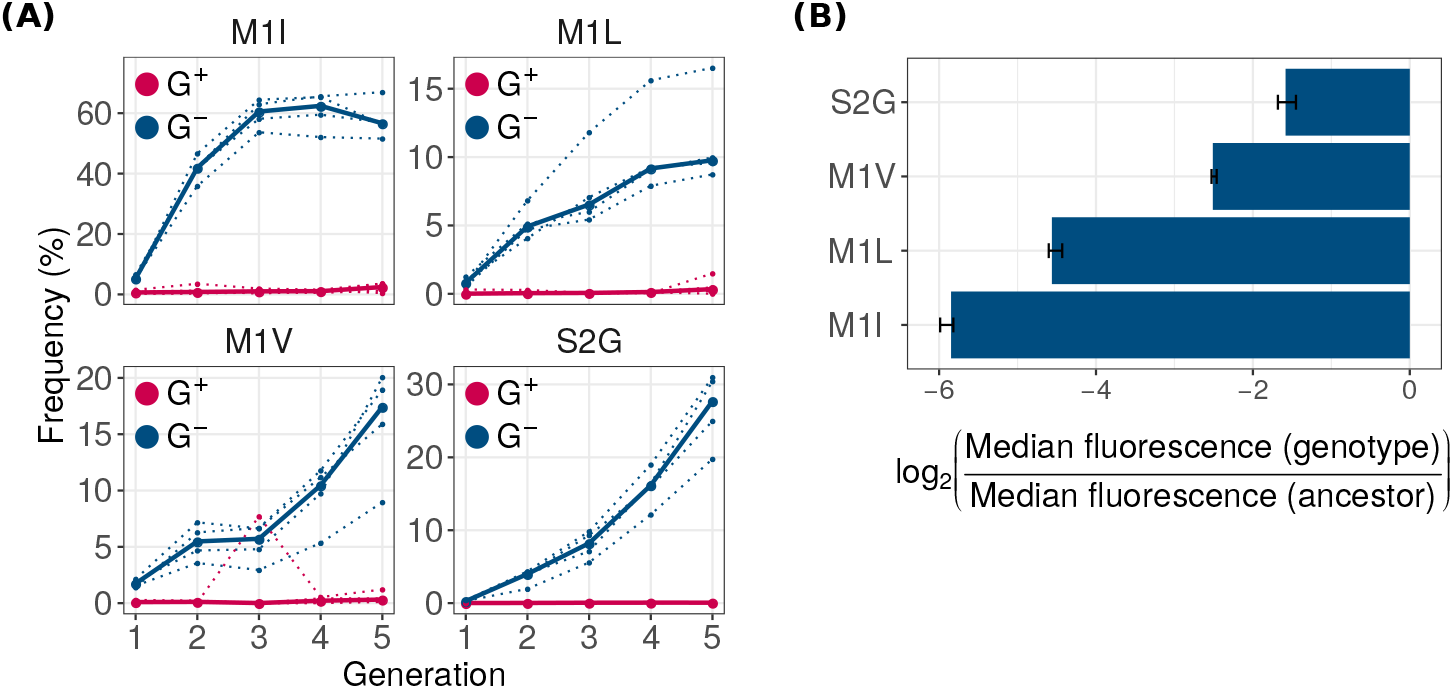
GroE expression disfavors accumulation of deleterious (fluorescence-reducing) mutations in phase 1. **(A)** Rise of deleterious mutations in evolving populations. The vertical axes show the frequency of deleterious mutations M1I, M1L, M1V and S2G in evolving G^+^ (red) and G^-^ (blue) populations during phase 1, at different generations (horizontal axes). The dotted lines denote the frequency of a mutation in individual replicate populations. The solid lines denote the median frequency over all replicates. **(B)** Effect of the mutations on green fluorescence. The horizontal axis shows the log_2_- transformed ratio of median green fluorescence for a given GFP variant (vertical axis) and ancestral GFP. A negative value denotes a deleterious effect whereas a positive value denotes a beneficial effect. The length of the bar denotes the median value of the log_2_-transformed fluorescence ratios in the three replicate measurements whereas the errorbar spans the range of minimum and maximum values.

To prove that these mutations indeed reduce fluorescence, we engineered them individually into the ancestral GFP using site directed mutagenesis, and measured their fluorescence. They caused a 2.7 to 64 fold reduction in median green fluorescence relative to ancestral GFP (***Figure 5B***), and are thus strongly deleterious to fluorescence. Their lower frequency in G^+^ populations demonstrates that GroE can potentiate the effects of individual deleterious mutations. Or experimental data on multi-mutant genotypes also shows that these mutations do not simply hitchhike to fixation with other, beneficial mutations (***SOM section 6***).

These observations raise the question why strongly fluorescence-deleterious mutations can become highly abundant in G^-^ populations in the first place. Since these mutations do not increase fitness by enhancing GFP activity, the likely reason is that they provide a growth advantage to cells harboring them. For example, three of these mutations (M1I, M1L, M1V) are start codon mutations. Such mutations can reduce the translation initiation rate^40^, the amount of synthesized protein, and hence also the protein expression cost^41^. Cells carrying these GFP mutations might have a lower metabolic burden and can outgrow other cells that synthesize more GFP^41^. To find out whether this is indeed the case, we measured the maximum growth rate of cells carrying the mutations M1I, M1L, M1V and S2G, relative to that of ancestral GFP (Materials and Methods), and found that these mutations indeed provide a significant growth advantage (Mann-Whitney U-test, *P* < 0.013). Thus, mutations that are deleterious for fluorescence can accumulate when GroE is not overexpressed. We note that our choice of weak selection is advantageous to detect strongly fluorescence-reducing mutations that are potentiated by GroE, because such mutations can persist only under weak selection.

### GroE expression increases phenotypic heterogeneity in fluorescence irrespective of the genotype

Next we asked why deleterious mutations may be disfavored under GroE expression. GroE might have an overall negative effect on fluorescence irrespective of the mutation, or it might affect strongly deleterious mutations differently from weakly deleterious mutations. To distinguish these possibilities, we measured the fluorescence of 15 differentially enriched mutations that we had engineered into ancestral GFP, and did so under GroE overexpression (***SOM Section 8***). We observed that for all mutants and for ancestral GFP, GroE overexpression caused the members of an isogenic population expressing a GFP variant to become increasingly heterogeneous in their fluorescence. (***Figure S14A***). Most strikingly, the distribution of the log-transformed fluorescence intensity became bimodal under GroE overexpression. One peak showed a higher and the other a lower fluorescence than the unimodal, Gaussian distribution 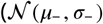 of fluorescence intensity without GroE overexpression.

The bimodal distribution of log transformed fluorescence intensity can be expressed as a sum of two Gaussian distributions 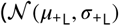 and 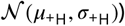, where the mean fluorescence at the lower peak (*μ*_+L_) and at the higher peak (*μ*_+H_) amount to an average of 93 and 107% of the mean log-fluorescence in the absence of GroE overexpression (*μ*_-_), respectively (***Figure S16, Table S3***). These results suggest that both buffering and potentiation are possible for the same GFP variant, depending on the cell where it is expressed.

### Phenotypic heterogeneity leads to the potentiation of some deleterious mutations but the buffering of others

To understand how this phenotypic heterogeneity may affect the selection of deleterious mutations, we developed a statistical model that relates fluorescence to fitness (i.e. the likelihood to survive experimental selection). We define the fitness of a genotype as the fraction of cells in an isogenic (genotypically homogeneous) population whose fluorescence intensity lies above the selection threshold we used in our directed evolution experiments. In the absence of GroE expression, individual cells of a given genotype show a unimodal Gaussian distribution with a mean *μ_-_* and variance *σ_-_* that we can estimate from our engineered mutants(***Figure S14***, ***Table S3***). In the presence of GroE expression, this distribution changes to a bimodal distribution whose parameters we can also estimate from data (***Figure S14***, **Table S3**). With this information in hand, we calculated the change in fitness of a genotype under GroE expression as the average difference in its fitness with and without GroE expression (ΔF = F_+_ - F_-_; Materials and Methods). A deleterious mutation with positive ΔF has a higher chance of surviving selection when GroE is expressed than when it is not. Thus GroE has a net buffering effect on this mutation. Conversely, a mutation with a negative ΔF has reduced chance of selection under GroE expression, and is hence potentiated by the chaperone.

Using this data-driven model, we found that GroE buffers strongly deleterious mutations whose fluorescence mean (*μ*_-_) lies no more than 5% above the threshold value that is needed for survival in our experiment. In contrast, GroE potentiated moderately deleterious mutations whose *μ*_-_ lies between 5 and 35.6% above this threshold (***Figure 6***). Outside this range, the value of ΔF is zero, and GroE neither buffers nor potentiates mutational effects.

**Figure 6.**
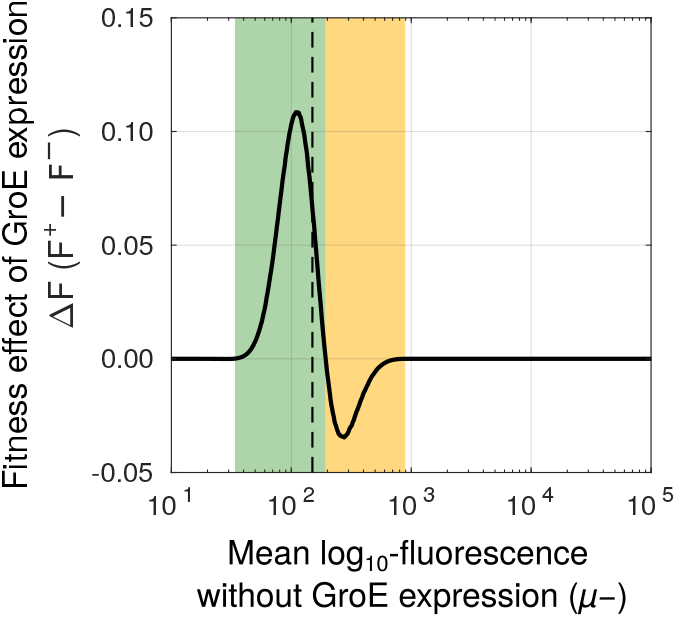
GroE buffers some mutations whereas it potentiates others. Change in fitness (ΔF, vertical axis) due to GroE expression predicted by our statistical model (Materials and Methods), as a function of mean log_10_ transformed fluorescence (***μ***-, horizontal axis) in the absence of GroE expression. The black dashed vertical line denotes the selection threshold (150 arbitrary units of fluorescence). A positive value of ΔF indicates buffering (green area) whereas a negative value indicates potentiation (orange area).

This model can explain several of our experimental observations, if one keeps in mind that our populations evolved under weak selection for fluorescence, and that individuals can accumulate deleterious mutations and survive selection as long as they fluoresce above a low fluorescence threshold. Even mutants whose mean fluorescence lies slightly below the threshold can persist at low frequency, because a few individuals may cross the selection threshold every generation due to phenotypic heterogeneity (***Figure S14***). Since most new mutations are deleterious^36,37^, fluorescence in our populations declines continually (***Figure 2***; G^-^ populations) until most genotypes fluoresce barely above the threshold. Our model predicts that GroE reduces the fitness of such deleterious but above-threshold genotypes, causing them to become depleted in G^+^ populations. This prediction is supported by our genetic diversity analysis (***Figure 3***). The model also predicts that mutations which are less deleterious and reduce fluorescence by a smaller amount, can persist in G^+^ populations (***SOM Section 8***), because GroE has no effect on the fitness of these mutations.

In addition, the model can help explain that some highly deleterious mutations become enriched in G^+^ populations, because GroE can buffer the deleterious effects of such mutations. One such mutation is a start-codon mutation M1T discussed in ***SOM Section 8***, which becomes enriched in G^+^ populations, even though its mean fluorescence lies 5 percent below the selection threshold.

### GroE leads to evolution of higher fluorescence intensity but lower color shift during directional selection towards a new phenotype

Since mutations that bring forth a new protein phenotype are often deleterious and destabilize a protein^7-10^, we also asked how GroE may affect the adaptive evolution of a new phenotype. We thus conducted a phase 2 of our evolution experiment, in which we selected for the new phenotype of cyan fluorescence. Since green and cyan fluorescence are correlated phenotypes (***Figure S5***), a green-fluorescing variant with high expression or stability could have a higher absolute cyan fluorescence than a cyan-fluorescing variant with low expression or stability. To avoid this problem, we selected cells whose cyan fluorescence increased relative to green fluorescence (***Figure S5***). Phase 2 started with populations from the end of phase 1 (round zero of phase 2). We subjected these populations, to five additional rounds of directed evolution towards cyan fluorescence.

After every generation of phase 2 evolution, we measured cyan and green fluorescence of 10^5^ cells from G^+^ and G^-^ populations, and observed that median cyan fluorescence significantly increased in all populations during phase 2 (linear model [LM]: ANOVA, *P* < 0.005; ***Figure S6A***) with a concomitant decrease in median green fluorescence (LM: ANOVA, *P* < 3.6 × 10^-4^; ***Figure S6B***). Thus, our populations can evolve increased cyan fluorescence.

Next, we asked if GroE expression influences the rate of evolution towards the new color. To this end, we compared the cyan fluorescence of G^+^ and G^-^ populations. During every generation (including the starting population derived from the end of phase 1), G^+^ populations had higher median cyan fluorescence than G^-^ populations (Mann-Whitney U test, *P* < 0.015; ***Figure S6A***).

We next asked whether the faster rate of cyan fluorescence in phase 2 G^+^ population originated during phase 2, or whether it might stem from the already higher fluorescence of the starting G^+^ populations from the end of phase 1 (***Figure S6A***). To find out, we normalized the fluorescence of the phase 2 starting populations to the same value for G^+^ and G^-^ populations. Specifically, we pooled the fluorescence values of individual replicates of the initial G^+^ populations (round zero), calculated the median fluorescence of this pooled population, and divided the absolute fluorescence values from each replicate population by this median. We proceeded analogously for the G^-^ populations, dividing their fluorescence by the median of the initial fluorescence values from a pool of all G^-^ populations. Next, we compared the rate of increase of this normalized fluorescence for both G^+^ and G^-^ populations with a linear model and found that GroE expression did not have a significant effect on this rate (***Figure 7A***). Moreover at the end of evolution, normalized cyan fluorescence was not significantly higher in G^+^ than in G^-^ populations. This analysis suggests that the difference between G^+^ and G^-^ populations during phase 2 may result from differences accumulated during phase 1. However, we also note that after generation one of phase 2, median cyan fluorescence increased more rapidly during every generation and remained somewhat higher in each of the last three generations (***Figure 7A***, **Figure S6A**).

**Figure 7.**
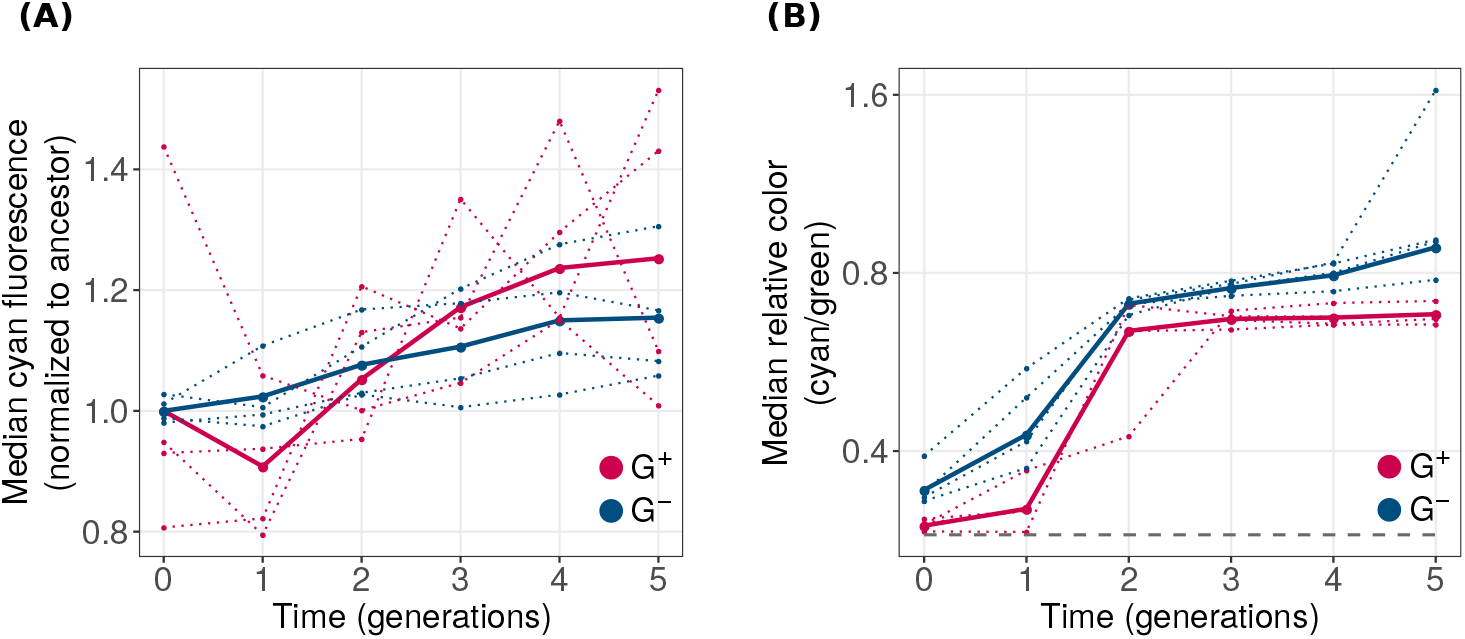
GroE expression leads to evolution of higher fluorescence but reduced color shift. Vertical axes denote **(A)** median normalized cyan fluorescence (normalized to starting populations; see main text), and **(B)** the relative color (cyan/green; logarithmically scaled) during phase 2 of directed evolution. The dotted line denotes the median value of individual replicates and the solid line denotes the median fluorescence value when data from all populations is pooled. Horizontal axes denote time in generations, where generation 0 refers to populations at the end of phase 1 evolution. G^+^ and G^-^ populations are indicated by red and blue colors respectively. The dashed horizontal line in panel **(B)** shows the relative color of ancestral GFP.

We also analyzed a different aspect of the phenotype, which is the extent of the spectral shift from green to cyan that occurred during phase 2. To find out whether GroE expression can affect the rate of this spectral shift, we calculated the ratio of cyan and green fluorescence for each cell in the different phase 2 populations. We refer to this ratio as relative color. Just like cyan fluorescence increased during phase 2 (***Figure 7A***), so did the spectral shift in both G^+^ and G^-^ populations (***Figure 7B***). However, this shift was lower for G^+^ populations than for G^-^ populations during every round of evolution (Mann-Whitney U test, *P* < 0.03).

### GroE reduces genetic diversity during evolution towards the new phenotype

We next asked whether GroE helps buffer deleterious mutations during phase 2, thus increasing a population’s genetic diversity, or whether it enhances their deleterious effects, thus reducing diversity. To find out, we sequenced the GFP coding region from each phase 2 population to an average coverage of 2155 sequences per population (750 - 3760 reads, depending on the population). Not surprisingly, the number of mutations per GFP coding sequence increased further during phase 2 evolution (LMM: ANOVA, *P* < 10^-10^; ***Figure 8A***), but we observed no significant effect of GroE expression on the rate of this increase (LMM: ANOVA, *P >* 0.05).However, when we quantified the genetic diversity of a population by the average pairwise distance between genotypes (***Figure 8B***), the diversity of G^+^ populations decreased during phase 2, whereas the diversity of the G^-^ further increased (LMM: ANOVA, *P* = 0.00015). Likewise, the Shannon entropy also significantly decreased in G^+^ populations compared to G^-^ populations (***Figure 8C***; LMM: ANOVA, *P* < 0.002). In sum, like in phase 1, GroE helps reduce genetic diversity, which is inconsistent with a net buffering of deleterious mutations, and supports the notion that GroE helps purge deleterious mutations. We also found that G^+^ populations had lower phenotypic diversity than G^-^ populations in every generation of phase 2 (Mann-Whitney U test, *P* < 0.0015). Just like in phase 1, principal component analysis shows that G^+^ populations accumulate different kinds of variants (***Figure S9***). Also, while potentiation dominates in its effect on genetic diversity, at least some buffering takes place as well (***Figure S7***).

**Figure 8.**
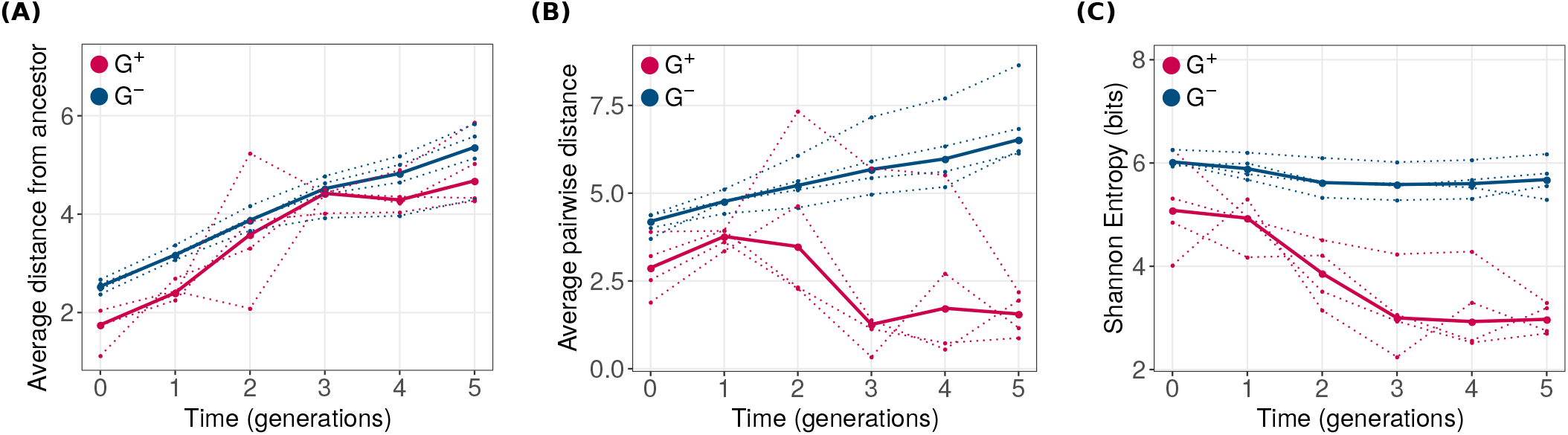
GroE expression leads to reduction of genetic diversity during phase 2 evolution. Genetic diversity metrics **(A)** average distance from the ancestral GFP, **(B)** average pairwise distance between genotypes and **(C)** Shannon entropy are shown on the vertical axes. Horizontal axes denote time in generations of evolution, where generation zero refers to the populations obtained after phase 1 evolution. G^+^ and G^-^ populations are color coded as red and blue, respectively. Average pairwise distance and Shannon entropy significantly reduced in G^+^ populations and were significantly lower than that of G^-^ populations (LMM: ANOVA, ***P*** < 0.002).

### GroE helps purge deleterious mutations during evolution of the new phenotype

G^+^ populations may acquire higher cyan fluorescence during phase 2 (***Figure 7A***) for two reasons. The first is that GroE may help spread mutations that convey the new phenotype. The observation that GroE delays evolutionary change in fluorescence color argues against this possibility (***Figure 7B**, **Figure S6C***). Adetailed analysis of specific mutants shows that it is indeed not the case (***SOM Section 9***).

The second possible reason is that GroE may help purge deleterious mutations from these populations, as it did during phase 1. If so, G^-^ populations should show lower fluorescence, because they preferentially accumulate deleterious mutations. To test this hypothesis, we identified single mutations that were significantly more abundant in G^-^ populations (GLM:LRT *P* < 0.05) relative to G^+^ populations by the end of phase 2. We restricted this analysis to variants whose frequency exceeded 5% at the end of evolution in at least one replicate population (***Figure S11***), and found 28 such mutations. Of these mutations, the most frequent were M1V, S2G and T203A. Each of them exceeded a frequency of 40% in every replicate population. We here focus on analyzing the mutations M1V and S2G. (***Figure S13***; see ***SOM Section 9*** for T203A). Both mutations are deleterious, reducing fluorescence (***Figure 5B***). Remarkably, they not only achieved a high frequency at the end of phase 2, but their frequency significantly increased during the five generations of phase 2 (generalized linear model with mixed effects: LRT *P* < 10^-15^). In contrast, their frequency did not increase in G^+^ populations where it remained below 0.5%. This suggests that these fluorescence-reducing mutations do not simply persist in G^-^ populations due to their higher abundance in the starting populations, i.e. the populations at the end of phase 1. GroE continues to help purge these mutations during selection for a new phenotype.

## Discussion

We used GFP as a model to find out whether GroE predominantly buffers or potentiates deleterious mutations that reduce fluorescence during protein evolution. Buffering refers to the suppression of a mutant’s deleterious effect. It occurs when a chaperone facilitates the folding of the mutant protein^18,23,24,27^. In contrast, potentiation refers to the enhancement of a mutation’s (deleterious or beneficial) effect^20,31,32^. We focused on deleterious mutations, because such mutations are most abundant during both stabilizing and directional selection^36,37,42^. If buffering is prevalent during stabilizing selection for an ancestral phenotype, then genetic diversity should increase over time, because mutations that are otherwise deleterious can accumulate in our evolving populations. In contrast, if potentiation is prevalent, genetic diversity should decrease, because deleterious mutations become eliminated more rapidly.

We found that GroE reduces genetic diversity during experimental evolution, implying that it helps purge deleterious mutations. It has been proven beyond doubt that GroE can buffer deleterious or destabilizing mutations^17,18,28,29^. Our experiments do not challenge this fact, because we show that GroE can indeed buffer some deleterious mutations. However, potentiation, not buffering is the dominant phenomenon in our evolution experiments. The notion that a chaperone can potentiate deleterious mutations is counter-intuitive, given that its protein folding assistance is expected to enhance protein function. However, it is not without precedent. For example, increasing the concentration of the chaperone Hsp70 can reduce a client protein’s folding yield^43^. We provide evidence that such potentiation is not just possible but can be dominant in the Hsp60 chaperone GroE.

During our evolution experiments, GroE reduced genetic diversity both under stabilizing selection for the ancestral green fluorescence phenotype, and under directional selection for a new (cyan) phenotype. During directional selection, it not only helped purge deleterious mutations, but also prevented the accumulation of key mutations with the new phenotype. Furthermore, GroE increased phenotypic heterogeneity in isogenic populations. That is, it enhanced fluorescence in a subset of a population, and decreased this activity in another subset. Thus GroE can buffer or potentiate the activity even of a single genotypic variant, depending on the cell in which the variant is expressed. The biochemical causes of this phenomenon remain to be determined. Overall, we find that GroE can buffer highly deleterious mutations whereas it potentiate mutations that are less deleterious.

One of few detailed studies on how GroE affects the evolution of individual proteins provided evidence for the importance of buffering^17^. The study found that during stabilizing selection for an enzyme’s ancestral phenotype, about 20-30% of enzyme variants that had evolved under GroE overexpression lost their activity when the chaperone was no longer expressed. In addition, GroE dependent variants evolved higher catalytic activity towards a novel substrate during directional selection. These experiments differed from ours in at least three ways that may help explain the prevalence of buffering. Firstly, they evolved enzymes. An enzyme’s activity depends not just on protein expression, folding and stability, but also on molecular motions that affect the rate of catalysis, whereas such motions play no role for our fluorescence phenotype. It is possible for a mutation to simultaneously enhance an enzyme’s catalytic activity and reduce its stability, an effect that is prominent among activity altering mutations in enzymes^18^. Since chaperones directly alter protein folding and stability but affect an enzyme’s catalysis only indirectly, it is possible to select mutations with such a stability-activity tradeoff. For a fluorescent protein there may be no such tradeoff, and variants with high activity (fluorescence) are also likely to be stable and thus chaperone-independent.

A second difference between our experiments and this previous work^17^ is that it used stringent selection where survival required catalytic activities to exceed 70% of the ancestral activity. In contrast, we deliberately used a relaxed selection strategy to expose chaperone effects. Finally, the previous work evolved small populations of approximately 200 variants, whereas we evolved large populations of *>* 10^5^ individuals. We were thus able to analyze the effects of GroE on a wider spectrum of variants.

Another relevant study^32^ focused on the effect of the chaperone Hsp90 on morphology-altering mutations in the yeast *Saccharomyces cerevisiae.* It defined potentiation as an increase, and buffering as a decrease in the variation of a morphometric trait caused by a chaperone. The study showed that Hsp90 mediated potentiation far outweighs buffering, except for mutations that have undergone several generations of selection under Hsp90 expression, which are buffered. This study supports our observation that chaperone mediated potentiation can be more important than buffering.

One limitation of our work is the assumption that different GFP variants do not affect the growth rate of the host. Such minimal interference with the host was one of the motivations to choose the non-native GFP, and to express it from a low-copy-number plasmid (Material and Methods). However, some of our start-codon mutations actually increased the host’s growth rate. To avoid this problem, future experiments might restrict mutagenesis to exclude such start-codon mutations. However, it may be difficult to eliminate growth-affecting mutations completely.

Our study opens exciting directions for future work. For example, the prevalence of potentiation or buffering may depend on the chaperone, the client protein, and on multiple other factors, such as selection strength and population size. More importantly, the observation that GroE induces phenotypic heterogeneity even among genetically identical cells calls for more detailed biochemical analysis of chaperone action. This unexpected complexity shows that studies on proteins amenable to single-cell phenotyping will be crucial to understand the mechanisms behind chaperone action and their role in adaptive evolution.

## Materials and Methods

### Construction of the expression system

#### Construction of the expression plasmid

We constructed a plasmid to express both GFP (constitutively) and GroE (inducibly). Our starting point for plasmid construction was the pGro7 plasmid designed by Takara^44^ for arabinose inducible expression of the chaperone proteins GroEL and GroES. This is a low-to-medium copy number plasmid with the pACYC origin of replication. It encodes chloramphenicol acetyltransferase, the transcription factor *araC* from *Salmonella typhimurium,* and the *groE* operon consisting of GroEL and GroES downstream of the *araBAD* promoter from *S. typhimurium.* Because we did not know whether leaky expression of pGro7 might occur even in the absence of arabinose, we created a control plasmid that cannot express the chaperone proteins at all. To this end, we digested pGro7 with BamHI and re-ligated the larger fragment corresponding to the plasmid backbone so as to eliminate the GroE operon. We named this control plasmid pΔGro7.

We next identified a region in pGro7 that can be used to place a GFP expression cassette. This region is a short stretch of DNA flanked by 5’-BglII and 3’-HindIII restriction sites downstream of the GroE operon. We use the GFPmut2 variant of GFP, which is distinguished by three amino acid changes from *Aqueoria vectoria* GFP^45^. This GFP variant is advantageous for our experiments because it is weakly dimerizing, has a single excitation peak (488nm), and undergoes fast maturation^46^. We obtained the GFP expression cassette, which consists of a promoter followed by a ribosome binding site and the GFP coding sequence, from plasmid pMSs201^47^. The GFP coding sequence is additionally flanked by 5’-XhoI and 3’-XbaI restriction sites. Since these sites already exist in pGro7 and are thus not useful for cloning, we engineered a 5’-SalI site and a 3’-SacI site flanking the GFP coding sequence in addition to the original restriction sites. We did so by PCR-amplifying the plasmid with the primers, pMS-Sal1-GFP-F and pMS-GFP-SacI-R (***Table S4***), and cloned the PCR-product back into the plasmid backbone. Next, we amplified the modified GFP expression cassette using the primers pMS-BglII-F and pMS-HindIII-R (***Table S4***), and cloned it into pGro7 and pΔGro7.

To identify the best promoters for GFP expression, we repeated this process with three variants of plasmid pMSs201, thus creating three pGro7 and three ΔGro7 plasmid variants that drive GFP expression from the ompA, rpsM and rplN promoters^47^. We quantified GFP expression from each promoter as explained in the next section.

#### Estimating of growth rates associated with different promoters

The host organism for our experiments is *E. coli* strain BW27784 (CGSC 7881), which cannot metabolize arabinose. We cultured all cells hosting our expression plasmids in LB with 25μg/ml chloramphenicol (LB+chl). Visual inspection of plated cells under blue light yielded green colonies and showed that all constructed plasmids expressed GFP. We corroborated this observation by measuring fluorescence on a plate reader (Tecan Spark 10M; ***Figure S1A***). To this end, we diluted 200μl of overnight (LB+chl) culture in 1ml PBS, distributed the diluted suspension into a 96 well plate in triplicate, and measured fluorescence in the GFP channel (485±10nm excitation, 521 ±10nm emission). Applying this procedure to each of our three pGro7 plasmids showed that GFP expression (fluorescence) from ompA and rplN promoters was 4.5 and 2.35 times higher than that from the rpsM promoter (***Figure S1A***), making these promoters better candidates for our experiments.

Next, we quantified the growth cost associated with GFP expression from the rplN and ompA promoters. To this end, we inoculated 30μl of overnight cultures that carried the corresponding pGro7-GFP plasmid in 14ml tubes containing 3ml LB+chl. After 60 minutes of growth at 37°C, we transferred 700μl of each culture to separate tubes and added different amounts of L-arabinose (from a 20% w/v stock solution) for GroE induction, such that the final arabinose concentrations equaled 0, 1 and 4 mg/ml. Next, we transferred 200μl from each culture to a 96-well plate (in triplicate). We measured optical density (OD at 600nm) and GFP fluorescence every 12 minutes during a growth period of 24 hours on a Tecan Spark 10M plate reader with temperature being maintained at 37°C, and with the plate shaken constantly between measurements. We inoculated and measured the growth of cultures with the two ΔGro7-GFP plasmids in the same manner. Using the final OD as an indicator of the carrying capacity, we fitted a logistic growth equation to the OD data using the fminsearch function (unconstrained, derivative free optimization) from the Optimization Toolbox in MATLAB (2017b), and estimated the growth rate from the fitted equation. Under arabinose induction, the growth rate was higher for the rplN promoter strain (***Figure S1B***) while the end point OD was comparable between the two promoter strains (***Figure S1C***). Therefore, we chose the pGro7-rplN-GFP (***Figure S2***) plasmid for all evolution experiments.

#### Measurement of GroE expression using SDS-PAGE

To determine the extent to which chaperone proteins are expressed from our plasmid at different concentrations of arabinose, we extracted total protein from the cells, performed SDS-PAGE of the protein extracts and observed the intensity of bands corresponding to proteins of the appropriate size. To this end, we first inoculated 30μl of overnight culture of the pGro7-rplN-GFP strain in 3ml of LB+chl and induced GroE expression with nine different concentrations of L-arabinose - 0, 0.002, 0.004, 0.008, 0.016, 0.04, 0.1,1 and 4 mg/ml - after 60 minutes of growth at 37°C. In these experiments, we also included the pΔGro7-rplN-GFP strain as an additional control for no plasmid-borne GroE expression. We allowed cell populations to grow for 8 hours. For each population, we pelleted cells from 1ml of cell suspension by centrifuging at 8000g for 3 minutes. We resuspended each pellet in 300μl of lysis buffer, which consists of 50mM Tris-HCl pH 7.5,100mM NaCl, 5% (v/v) glycerol, 1mM dithiothreitol (added fresh), 1x protein inhibitor cocktail (cOmplete, Roche; added fresh), 300μg/ml lysozyme, 3μg/ml DNAseI, and 16mM MgCl_2_. We then incubated this suspension for 4 hours at 4°C. We lysed the cells by freezing the suspension in liquid nitrogen, followed by thawing it in a water bath, and repeated this freeze-thaw cycle ten times. Then, we centrifuged the suspension at 18000g for 30 minutes at 4°C and collected the supernatant. We quantified protein concentration using the Bradford method (Bio-Rad Quick Start™ Bradford reagent). We then heated 10μg of protein sample with suitable amounts of 4x SDS-PAGE loading buffer (250 mm Tris-HCl pH 6.8, 8% w/v SDS, 0.2% w/v bromophenol blue, 40% v/v glycerol and 20% v/v 2- mercaptoethanol) at 95°C for 5 minutes. We loaded the samples on a polyacrylamide gel (4% for stacking and 12% for resolving; TruPAGE precast gel, Sigma-Aldrich), and performed electrophoresis at 180V for 45min in 1x TruPAGE TEA-Tricine SDS buffer (Sigma-Aldrich). We fixed the gel for 30 minutes in fixing/destaining solution (50% v/v methanol, 10% v/v acetic acid), and stained it overnight in Coomassie brilliant blue staining solution (0.1% w/v Coomassie brilliant blue R-250, 50% v/v methanol, 10% v/v acetic acid). Next, we destained the gel with destaining solution until the background was clean and the bands were clear.

We observed no induction of GroEL (60Kda) in the absence of arabinose (***Figure S3***) but strong induction even at the lowest tested concentration of arabinose (0.002μg/ml). The 60Kda GroEL band was missing in both the pΔGro7-rplN-GFP sample and in the no-induction sample, suggesting that leaky expression is negligible. With these observations in mind, we chose a modest concentration of 0.1mg/ml arabinose for induction in all subsequent experiments. We reasoned that at this concentration of arabinose the expression of GroE would be saturated, and small deviations from this chosen value during the experiments would not affect the expression.

### Mutagenesis and selection

#### Preparation of electrocompetent cells

To prepare electrocompetent cells, we performed every step of the procedure described below in detergent-free glassware. We cultured *E.coli* strain BW27784 in 10ml SOB medium overnight at 37°C with shaking at 220rpm. Subsequently, we inoculated 1l of pre-warmed (37°C) SOB in a 5l flask with the overnight culture. We let cells grow for 2 - 3 hours (37°C+220rpm) until their OD reached 0.4-0.6. Then we transferred the flask to ice and let it cool for 20 minutes. Subsequently, we transferred the cell suspension to two 500ml centrifuge bottles (Eppendorf), and centrifuged both bottles at 1500rpm for 15 minutes at 4°C with neither acceleration nor deceleration, on a swinging bucket rotor (Eppendorf). Next, we resuspended the cells in 90ml of cold water per bottle by gently swirling the bottle, and distributed the suspension in six pre-chilled 50ml tubes (30ml per tube). We gently added 15ml of cold glycerol-mannitol solution (20% w/v glycerol, 1.5% w/v mannitol) to the bottom of each tube. Then we centrifuged the tubes at 1500g for 15 minutes at 4°C without acceleration/deceleration. We removed the supernatant and resuspended the pellet of each tube in 1.5ml of cold glycerol-mannitol solution. We combined the cell suspension from all tubes and aliquoted 100μl into chilled 1.5ml tubes. We snap-froze aliquots in liquid nitrogen bath and stored them at −80°C.

#### Mutagenesis

For mutagenesis by error-prone PCR, we used the primers Gro-Mut-F and Gro-Mut-R to amplify GFP from pGro7-rplN-GFP (***Table S4***).

For the error-prone PCR itself, we used the following reaction mixture: 150nM each of the nucleotide analogs 8-oxodeoxyguanosine triphosphate (8-oxo-dGTP, Trilink Biotechnologies) and 6-(2-deoxy-beta-D-ribofuranosyl)-3,4-dihydro-8H-pyrimido-[4,5-C] [1,2]oxazin-7-one triphosphate (dPTP, Trilink Biotechnologies), 200nM each of forward and reverse primers, 400nM of each dNTP (Thermo Scientific), 1x ThermoPol buffer (NEB) and 25 units/ml of Taq polymerase (NEB). We prepared 100μl of the PCR reaction with 5ng of plasmid DNA as the template (~6×10^8^ molecules), and split the reaction mixture into two 50μl aliquots for efficient heat transfer during PCR. We performed the PCR with the following program: Initial denaturation at 95°C for 5 minutes, 25 cycles of amplification with 95°C for 30 seconds, 56°C for 30 seconds, 72°C for 1 minute, and a final extension at 72°C for 5 minutes. We optimized this protocol such that we obtained ~1 nucleotide mutation per amplicon corresponding to approximately 0.95 amino acid changes per GFP protein.

We purified PCR products using a QIAquick PCR purification kit (QIAGEN). Subsequently, we prepared 50μl of restriction digestion reaction with 400ng PCR product, 20 units each of the two restriction enzymes, SalI-HF and SacI-HF (NEB), 20 units of DpnI (NEB; for removing template plasmid), and 5μl of 10x-CutSmart buffer (NEB). We carried out the digestion overnight and purified the digested products with the QIAquick PCR purification kit. We digested the plasmid in the same way using the restriction enzymes. Additionally, we treated the plasmids with Antarctic phosphatase (NEB) to minimize their self-ligation. We separated the digested plasmid backbones from the insert using agarose gel electrophoresis, and purified the excised band using a QIAquick gel extraction kit (QIAGEN). For ligation, we prepared a 30μl ligation mixture consisting of 100ng of the digested plasmid backbone, 55ng of the digested amplicons (1:3 molar ratio of backbone and insert), 3μl of 10xT4 DNA ligase buffer (NEB), and 1.5μl (600 units) of T4 DNA ligase (NEB). We performed the ligation overnight at 16°C. To separate salts from the ligation products, we added 70μl water, 50μg glycogen (Thermo-Fisher), 50μl 7.5M ammonium acetate and 375μl ethanol to the ligation mix. After incubating the mixture for 20 minutes at −80°C, we centrifuged it at 18000g for 20 minutes at 4°C. We decanted the supernatant and washed the pellet twice with 800μl of 70% ethanol. We dried the pellet and resuspended it in 20μl of sterile deionized water.

#### Transformation of the mutant library using electroporation

We thawed frozen electrocompetent cells on ice and added the purified ligation products to them. We transferred the resulting suspension into a 2mm electroporation cuvette (EP202, Cell Projects, UK), and performed electroporation with a single 3kV pulse using the Bio-Rad MicroPulser (program EC3). We immediately added 1ml of warm (37°C) SOC medium, transferred the suspension to a 35ml glass tube (17mm diameter), and incubated the cells for 1.5 hours at 37°C with shaking at 220rpm. We plated 100μl of a 512-fold diluted suspension (three 1:8 serial dilution) on an LB-chl agar plate and added 9ml of LB-chl to the undiluted suspension. We incubated the plates and the tubes (with shaking at 220rpm) overnight at 37°C. We estimated the library size by counting the number of colonies on the LB-chl plate. Throughout our evolution experiments, we maintained a minimum library size of 10^5^ transformants.

#### Estimation of mutation rate

To estimate the mutation rate of our mutagenesis procedure, we performed mutagenesis on the ancestral GFP gene and transformed the mutants using electroporation as described in the previous section. We performed colony PCR with ten randomly picked colonies from the plate and sequenced the PCR products using Sanger sequencing to estimate the mutation rate. In this way, we determined the mutation rate of the ancestral GFP gene during every round of directed evolution to ensure that it stayed in the range of 1-2 mutations per amplicon throughout the evolution experiment. It is well-known that PCR-mutagenesis creates a biased mutation spectrum^39^, and our protocol is no exception. From the combined Sanger sequencing data obtained from all rounds of evolution, we estimated the frequencies of different point substitutions as follows: AT→GC: 0.755, GC→AT: 0.144, AT→TA: 0.072, AT→CG: 0.025, GC→CG: 0.004 and GC→TA: 0. Thus, the protocol is biased towards AT→GC transitions.

#### Selection of transformed cells using FACS

We performed directed evolution in four replicate populations where GroE was expressed from our expression plasmid, along with four control populations in which it was not expressed from this plasmid. We applied the following selection protocol to each population. To prepare for selection, we inoculated 4ml of LB-chl in a 20ml glass tube with 80μl of the appropriate transformed library. We allowed the cells to grow at 37°C with shaking at 220rpm for 60 minutes, and then induced GroE expression in G^+^ populations with 0.1 μg/ml of L-arabinose. We allowed cells to continue their growth for another 10 hours. Subsequently, we transferred the tubes to ice and pelleted cells from 700μl of the suspension by centrifuging at 8000g for 3 minutes. We washed cells by resuspending them in cold PBS and centrifuging them again. We decanted the supernatant, resuspended the cells in 1ml cold PBS, and transferred 100μl of the suspension to 1ml cold PBS in a 5ml polystyrene tube (Falcon). We performed cell sorting on a BD FACSAriaIII cell sorter with the following photomultiplier tube (PMT) voltages for different channels - 478V for FSC, 282V for SSC, 480V for FITC and 493V for AmCyan. We excluded debris and other small particles by setting a threshold of 1000 on FSC-H and SSC-H.

We used the FITC channel (488nm excitation and 530±15nm emission) for measuring green fluorescence and the AmCyan channel (405nm excitation and 510±25nm emission) for measuring cyan fluorescence. We quantified the autofluorescence of cells in each channel by measuring the fluorescence of untransformed cells. To select variants with green fluorescence, we sorted cells with a FITC-H value higher than the maximum FITC-H value of the untransformed cells. Because green and cyan fluorescence are correlated - wild-type GFP fluoresces in both the FITC (green) and the AmCyan (cyan) channel - we did not define the new phenotype merely as a higher fluorescence in the AmCyan channel. Instead, we required a relative shift towards cyan fluorescence that cannot be solely explained by higher green fluorescence. Specifically, we plotted the fluorescence of wild type GFP in the two channels (FITC-H and AmCyan-H) against each other, and designated the area that lay both above the regression line and the background fluorescence of the AmCyan channel as the selection gate (***Figure S5***). This procedure ensures that surviving cells show cyan fluorescence that cannot be merely explained by enhanced green fluorescence.

We sorted 10^5^ cells into 1.5ml tubes containing 500μl of cold LB. We incubated the sorted cells at 37°C for 30 minutes and then transferred them to 5ml of LB-chl in a 20ml glass tube. We let the cells grow overnight at 37°C with shaking at 220rpm. We inoculated 4ml of LB-chl with 80μl of the overnight culture and repeated the induction and the sorting procedure as described above. We performed the second sort to minimize possible contamination from cells that did not meet our selection criteria. We incubated the sorted cells at 37°C for 30 minutes and transferred them to 10ml LB-chl in a 50ml tube. We allowed these cells to grow overnight and used 1ml of the overnight culture for preparing glycerol stocks (15% glycerol). We used the remainder of the culture for extracting the plasmid library using a QIAprep Spin Miniprep kit (QIAGEN). The plasmid libraries thus isolated served as templates for the next round of mutagenesis.

### Analysis of fluorescence of populations using flow cytometry

We used flow cytometry to analyze the phenotype of evolving populations after every generation, i.e., after every round of mutagenesis and selection. To this end, we first obtained an overnight culture either directly after the second round of sorting (previous section), or by reviving a glycerol stock. From this culture, we inoculated 40μl of cell suspension in 4ml LB+chl. After 1 hour of growth at 37°C with shaking at 220rpm, we added L-arabinose to a final concentration of 0.1 μg/ml, and allowed the cells to grow for another 9 hours. Next, we pelletted cells from 500μl of the culture by centrifuging at 8000g for 3 minutes at 4°C. Then, we washed the cells by resuspending them in 1ml cold PBS and pelletted them again. We resuspended the cells in 1ml PBS and transferred 60μl of this suspension into 1ml of cold PBS in a 5ml polystyrene tube (Falcon). We quantified green fluorescence using the FITC channel (488nm excitation and 530±15nm emission), and cyan fluorescence using the AmCyan channel (405nm excitation and 510±25nm emission) on a BD LSR FortessaII flow cytometer. The PMT voltages for the FITC and AmCyan channels were 480V and 493V, respectively. We recorded 100,000 events and analyzed the data using both MATLAB (fca-Readfcs.m^48^) and the R package flowCore^49^. We note that GroE expression led to an increase in number of non-fluorescent “events” (signals) even in an isogenic population (data not shown). We surmise that these non-fluorescent events could originate from nonviable cells which in turn could arise due to protein overexpression stress. Therefore, we excluded all non-fluorescent cells from our analyses.

We measured the fluorescence of evolved populations after every round of directed evolution. To analyze the temporal change in fluorescence we fitted a linear mixed model using the R package lme4(v1.1-21)^50^. In this analysis, we used fluorescence (green in phase 1 and cyan in phase 2) as the response variable, with time (round of evolution) and state of GroE expression (condition: G^+^ or G^-^) as interacting predictors, and variation between replicates as random effects. Specifically, we used the following expression to define our statistical model: `Fluorescence - rounds*condition + (1|replicate)`. Because all phase 1 populations started with the same ancestral fluorescence, we forced a constant intercept for the linear model. We analyzed the significance of the fit using the `anova` function from the R package lmerTest(v3.1-0)^51^. This function performs a type III analysis of variance of the fitted coefficients and estimates degrees of freedom using Satterthwaite’s method^51^. In the model we used, the factors (rounds of evolution and GroE expression) do not significantly affect the response variable (fluorescence), under the null hypothesis.

### Analysis of DNA sequencing data

#### Preparation of sequencing libraries

We sequenced the GFP coding sequence from plasmid libraries isolated after every round of evolution using single molecule real time (SMRT) sequencing(Pacific Biosciences, PacBio).

For multiplexed sequencing, we generated barcoded libraries according to the instructions provided by PacBio^35^. To create these libraries, we performed two rounds of PCR. In the first round we amplified the GFP coding region with primers carrying a “universal” sequence provided by PacBio. We amplified the GFP gene from the plasmid libraries obtained after every round of selection (see Selection of transformed cells using FACS), using the primers GLG_ORF_PacBio-F and GLG_ORF_PacBio-R (***Table S4***). These primers have a 5’ amino-C6 modification and had been PAGE purified before use. We prepared a 25μl PCR mix with 5ng of plasmid, 400nM of each primer, 200nM of each dNTP, 5μl of 5x Phusion HF buffer (Thermo-Fisher), and 0.5 units of Phusion High-Fidelity polymerase (Thermo-Fisher). We then performed PCR using an initial denaturation at 95°C for 5min followed by 20 cycles of amplification with the following program: 95°C for 30s; 58°C for 30s, 72°C for 30s, and a final extension at 72°C for 2min. We added 8μl of water, 1μl of 10x-CutSmart buffer (NEB), 10 units each of DpnI (NEB) and ExoI (NEB) to the PCR products, and incubated the mixture for 30 minutes at 37°C, followed by 10 minutes at 85°C. After the reaction, we diluted the mixture by adding 70μl of water.

We carried out a second round of PCR to barcode the different samples. We selected 36 barcode sequences from a list of 384 16nt-barcode sequences from PacBio^52^, and designed 18 barcoded forward and reverse primers. All these primers carry a 5’ phosphate modification and were HPLC purified before use. For this second PCR, we used 2μl of the DpnI-ExoI treated first-round PCR product (diluted) as the template in a 50μl PCR mix consisting of 200nM of each of the barcoded primers, 200nM of each dNTP, 10μl of 5x Phusion HF buffer (Thermo-Fisher), and 1 unit of Phusion High-Fidelity polymerase (Thermo-Fisher). We performed PCR using an initial denaturation at 95°C for 5min, followed by 30 cycles of amplification using the following program: 95°C for 30s, 62°C for 30s, and 72°C for 40s, and a final extension at 72°C for 2min. We determined the purity of the PCR products through agarose gel electrophoresis, and found that most of the products were clean, without non-specific bands or primer dimers. We purified these products using a QIAquick PCR purification kit (QIAGEN). For the few samples that contained large amounts of primer dimers, we purified the PCR products using gel extraction. We measured the concentration of the purified products using a Nanodrop 1000 (Thermo-Fisher) spectrophotometer and a Qubit 3.0 fluorometer (Life Technologies). For GFP libraries from each phase of evolution (1 and 2), we pooled 130ng of every barcoded product and purified the two resulting pools using gel extraction.

For subsequent DNA sequencing two PacBio Sequel SMRT cells were used for the two pooled samples (phase 1 and phase 2), and sequencing was performed on the PacBio Sequel system (3.0 Chemistry) by the Functional Genomics Center Zurich.

#### Processing of raw data

We obtained approximately 600,000 raw zero mode waveguide (ZMW) reads from each of the SMRT cells. We determined the consensus of circular sequences (CCS) from the raw data (subreads) using the ccs (v3.4.1) application from the PacBio SMRTlink package^53^, with the following parameters: minimum length = 750, maximum length = 1500, minimum passes = 3, minimum predicted accuracy = 99%. We kept the other parameters at their default value. We obtained approximately 55% of the ZMW reads. We demultiplexed the post-CCS reads using lima (v1.9.0, SMRTlink), and aligned them to the reference sequence (GFPmut2) using minimap2 (v2.15-r905, SMRTLink). We analyzed the alignments in SAM format using a custom awk script. Since our sequencing data contained many indels, we confirmed that these were sequencing artifacts by Sanger-sequencing twenty random single clones, and excluded any indels from further analysis. From the data thus filtered, we obtained lists of single mutations as well as genotypes with multiple mutations, along with their raw counts and frequencies.

Synonymous mutations can alter co-translational folding^54^, but GroE-assisted folding occurs post-translationally. For this reason, we restricted our analysis to non-synonymous mutations.

#### Estimation of genetic diversity

We used three measures of genetic diversity, all of which are based on the observed number of amino acid changes in our evolving GFP sequences. The first is the average distance of the genotypes in a population from the ancestor, defined as the average number of amino acid mutations in a population relative to the ancestral GFP sequence. The second is the average pairwise distance between two genotypes in a population, defined as the Hamming distance between their amino acid sequences. This metric is analogous to a widely used nucleotide diversity metric^55^, except that we apply it to amino acid sequences. Specifically, we define

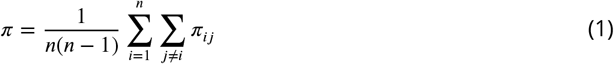

where *π* denotes the average pairwise distance between any two genotypes in a population, *π_ij_* denotes the distance between the *i^th^* and the *j^,h^* genotypes, and *n* is the total number of genotypes in the population.

The third metric is the Shannon entropy *H* of individual allele frequencies *p_i_* in a population^56^, defined as

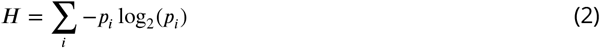

This diversity measure is largest if all alleles have equal frequencies, and it decreases as an allele frequency distribution becomes increasingly peaked at one or few alleles that occur at much higher frequencies than the other alleles.

We used linear mixed-effects models (LMM) implemented in the R package lme4 (v1.1-21)^50^ to analyze the effect of GroE overexpression on changes in genetic diversity between different populations. We represented diversity as the response variable, with time (rounds of evolution) and state of GroE expression (G^+^ or G^+^) as interacting predictor variables. We include differences between the replicate populations as random effects. Specifically, we used the following expression to define the model: `diversity - rounds*condition + (1|replicate)`. We tested the model’s goodness-of-fit to the sequence data using the `anova` function from the R package, lmerTest (v3.1- 0)^51^.

#### Principal component analysis of the genotypes

We performed principal component analysis (PCA)^39^ to visualize different genotypes accumulated in a population. To this end, we randomly sampled 200 sequences without replacement from every replicate population after the end of the final round of evolution. We converted the amino-acid sequence of each genotype to a numerical sequence, assigning a numerical code to each amino acid. Specifically, we assigned the numbers 1 to 20 to amino acids in the following order: W, F, Y, I, V, L, M, C, D, E, G, A, P, H, K, R, S, T, N, and Q. This ordering of amino acid, in contrast to an alphabetical order, ensures that chemically similar amino acids have a small numerical difference between them^57^. We assigned the number −10 to the stop codon, because effects of nonsense mutations are dramatically different from those of missense mutations.

We then performed PCA on a matrix containing all these numerical sequences, using the `prcomp` function from the R package stats (v3.4.4)^58^ (Figures S8A & S9A). The rows of this matrix harbor individual sequences (genotypes). Its columns correspond to individual positions in the sequence.

We also performed PCA on a matrix harboring allele frequencies of all single amino acid mutations from each population at the end of evolution (Figures S8B & S9B). In this matrix, each row contains allele frequency data from a different population, and each column corresponds to a different observed mutant.

#### Monte-Carlo simulations to calculate variant frequencies expected under mutation pressure alone

We restricted this analysis to amino-acid variants that attain a minimal threshold frequency in at least one replicate population at the end of evolution. We chose this threshold frequency to be 3.5% for phase 1, and 5% for phase 2 to keep the number of variants manageable for all subsequent analyses. No individual population had more than 56 variants exceeding these thresholds. For each of these variants, we performed Monte Carlo simulations to test the null hypothesis that mutation pressure alone may be responsible for the variant’s frequency. Rejection of this null hypothesis for any one variant implies that selection must be involved in explaining its frequency. (Our experimental populations are sufficiently large that genetic drift is negligible on the time scales of our experiment.)

This numerical analysis consists of two parts. In the first part, we compute the probability *π* that a specific amino acid variant arises in the population. In the second part, the Monte Carlo simulation proper, we simulate how the frequency of this variant changes over time due to mutation pressure alone.

We explain this procedure with the mutation S147P, which occurs at a frequency of less than 0.4% in all the G^+^ populations at the final (5th) round of phase 1. After five additional rounds of evolution in phase 2 this mutation attained a frequency greater than 98% in all replicate populations. S147P is encoded by the codon CCA, which requires a T→C change at the 439th position of the GFP coding sequence, which corresponds to the first position of the ancestral codon 147 (TCA→CCA).

For an S147P mutation to occur, three events must take place. We calculate their probability as follows.

- At least one mutation must occur (somewhere) in the GFP coding sequence. Because mutations are rare, we model the probability *P*_mut_ of this event with a Poisson distribution, such that

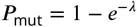 Here *λ* denotes the average number of mutations in each individual per round of mutagenesis. Because we had calibrated our mutagenesis protocol such that this number lies between 1 and 2 (see Estimation of mutation rate), we use a value of *λ* = 1.5, which leads to *P*_mut_ = 0.777
- One of the occurring mutations must affect the 439th nucleotide position, whereas all other mutations must occur outside codon 147. If only one mutation occurs in the GFP coding sequence, the probability of this event (*P*_pos_) is equal to the probability of choosing this one position from the GFP coding sequence of length 717nt, which is 1/717 = 0.0014. If two mutations occur in the coding sequence then *P*_pos_ would be 1/717 × 715/716 = 0.001393. If three mutations occur, *P*_pos_ = 1/717 × 715/716 × 714/717 = 0.001391. Analogous expressions apply for (increasingly unlikely) higher numbers of mutations. Since the difference between the values of *P*_pos_ for the above three cases is very small, we can conveniently approximate its value to be *P*_pos_ = 1/717 = 0.0014. Because this value is a slight overestimate, our statistical inference would be conservative.
- This mutation must cause a T→C change. From our estimation of mutation rates by Sanger sequencing experiments (Estimation of mutation rate), we know that this probability (*P*_sub_) is 0.75

The probability (*π*) that the mutation S147P occurs (i.e. all the above-mentioned events occur) in any one generation is the product of the above three probabilities: *P*_mut_ × *P*_pos_ × *P*_sub_ = 8.18 × 10^-4^.

We note that we can neglect amino acid changes caused by double or triple nucleotide mutations, because our sequencing data showed that every amino acid variant that exceeded our threshold frequency was caused by a single nucleotide change.

We next turn to the second part of our numerical analysis, where we use the probability *π* that a specific variant arises to calculate how the expected mean frequency of this variant changes over time. To this end, we used a discrete time stochastic model of a population whose individuals mutate at a rate *π*, such that the number of unmutated individuals becomes progressively smaller. Our simulations neglect back-mutations to the wild-type allele, which will slightly overestimate the allele frequencies caused by mutation pressure. In consequence, our analysis below will be statistically conservative. That is, it might accept some variants as having a frequency consistent with mutation pressure alone, while they may actually be affected by selection.

Specifically, for each mutant whose frequency exceeded our threshold, we performed the following simulation 10^5^ times, with a starting population of *N_0_* = 10^5^) individuals. Each individual has a probability *π* (as described earlier) of acquiring a given mutation. The number of individuals mutated in the first round of evolution is thus given by a random variable that is binomially *(**P**(π. N_0_))* distributed. In our simulations, we generated a pseudorandom number *M_0_* from this distribution, and computed the number of unmutated individuals after the first round of evolution as *N_1_* = *N_0_ - M_0_*. In the second round, the number of individuals experiencing the mutation is a random variable with binomial distribution, *B(π,N*_1_). We also generated an instance *M*_1_ of this random variate numerically, and calculated the number of unmutated individuals after the second round as *N_2_* = *N*_1_ - *M*_1_. We repeated this procedure for three more rounds/generations to obtain the frequency of the remaining wild-type alleles, and thus also the frequency of the mutant alleles at the end of phase 1. We repeated the procedure for an additional five rounds to obtain mutant allele frequencies at the end of phase 2 evolution. For each mutation, we performed 10^5^ such simulations and calculated the fraction of simulations in which the predicted frequency exceeded the threshold frequency of 3.5% for phase 1 and of 5% for phase 2.

For each variant whose frequency exceeded the threshold in our experimental populations, not a single one among 10^5^ simulation reached this threshold. Thus, if we consider the null-hypothesis that the observed frequency of any one variant can be explained by mutation pressure alone, our simulations reject this null-hypothesis at a p-value of *P* < 10^-5^. Applying a Bonferroni correction to the number of such tests we performed (<90 tests, i.e., variants, for each of the two threshold), we reject the null-hypothesis at a Bonferroni-corrected P-value of *P* < 0.0009. In sum, the frequency of no mutation we consider here can be explained by mutation pressure alone. Since our populations are so large that we can neglect genetic drift at the time scale of this experiment, selection or hitchhiking with another high frequency mutation must be invoked to explain their frequency.

#### Calculation of mutation enrichment

For each round of evolution, we compared the enrichment of mutations in G^+^ populations relative to G^-^ populations, using generalized linear models (GLM; R stats package^58^ v3.4.4). Specifically, we fitted a GLM with a logit link function (binomial model) using mutation counts as the response variable, and the state of GroE expression (G^+^ or G^-^) as the predictor variable. We analyzed the goodness of fit of the full model (slope + intercept) with respect to a reduced (intercept only) model, using the `anova` function from the R stats package v3.4.4. This function performs an analysis of deviance on the models, using a likelihood ratio test, and determines if the additional parameters (slope in our case) significantly improve the fit. Here, a positive value of the slope denotes enrichment of the mutation in G^+^ populations. We adjusted the p-values thus obtained for multiple testing using a Bonferroni correction. We used the mutations with a corrected p-value of *P* < 0.05 in subsequent analyses.

#### Estimating the strength of selection acting on mutations

For the variants that satisfied the frequency threshold criterion of Monte-Carlo simulations, we estimated the strength of selection in G^+^ and G^-^ populations by fitting generalized linear mixed-effects models (GLMM) using the function `glmer` from the R package lme4 (v1.1-21)^50^. We fitted two models, one each for GroE overexpression and control populations, using a logit link function with mutation counts as the response variable, time (round of evolution) as the predictor variable, and the variation between replicate populations as the random effect. Specifically, we defined the models with the following expression: `Counts - rounds + (1|replicate)`. We analyzed the goodness of fit of the full model (slope + intercept) with respect to a reduced (intercept only) model, using the `anova` function from the R stats package v3.4.4. We adjusted the p-values thus obtained for multiple testing using a Bonferroni correction. The estimated value of the slope denotes the strength and direction of selection. A positive value denotes positive selection, and a higher absolute value of the slope denotes stronger selection.

### Construction and analysis of specific mutants

#### Engineering specific mutations

We used PCR-based site directed mutagenesis to engineer specific mutations into the GFP gene. To this end, we first created a “minimal” plasmid that expresses GFP constitutively (pMini-GFP) from the rplN promoter, but that did not contain the chaperone genes and the *araC* gene. We designed primer pairs to amplify this entire plasmid from the site of the desired mutation (***Table S5***). Specifically, we designed these primer pairs with 15 complementary nucleotides at their 3’ end and a non-complementary region that did not exceed 25 nucleotides in length. We included the desired mutation in the complementary region. Whenever the difference in melting temperature (Tm) of the primers exceeded 5°C, we trimmed the non-complementary region of the primer with higher Tm from the 5’ end. We used the software tool — `melting` (ver. 5.1)^59^ to calculate Tm. We designed the primers in this way to minimize inefficient amplification due to primer dimer formation.

We amplified pMini with different primer pairs for each mutation to be engineered (***Table S5***) using high fidelity Q5 polymerase (NEB). We transformed the PCR products into *E.coli* BW27784 cells made transformation-competent with the CaCl_2_ method^60^, using a standard heat shock transformation method^60^. We isolated and purified plasmid from the clones thus obtained and sequenced the GFP gene to confirm the mutation. Next, we cloned each mutated GFP gene into the GroE expression plasmid, pGro7-rplN-GFP.

We generated double mutants via the same procedure, by engineering the mutations serially via two rounds of PCR. Next, we cloned the mutated GFP gene into pGro7-rplN-GFP.

#### Measurement of growth rates associated with different GFP mutants

To measure whether selected mutations (M1I, M1L, M1T, M1V and S2G) confer a growth advantage in the absence of GroE expression, we prepared 1:20 dilutions of an overnight culture of each mutant, as well as of the strain expressing ancestral GFP. We inoculated 10μl of each diluted suspension into 1.4 ml of fresh LB+Chl, and aliquoted 200μl of this inoculated medium into six wells (replicates) of a 96 well plate. Next, we measured optical density (OD at 600nm) every 10 minutes during 20 hours on a Tecan Spark plate reader at 37°C and with the plate shaken constantly between measurements.

All mutants reached stationary phase after 13 hours of growth. Because the growth data fitted a logistic growth equation poorly (not shown), we calculated the maximum growth rate, another commonly used estimate of growth^61^. To this end, we first logarithmically (base 2) transformed the measured OD values for each mutant. Using a sliding window of six time points (corresponding to 1h of growth), we calculated the rate of change of log_2_-OD between consecutive time points using a linear model (R stats package v3.4.4)^58^. Next, we calculated the maximum value of this slope for all time points, and for each of the six replicates for each mutant and the ancestor. We then compared the median maximum growth rate (of the six replicates) for each mutant to that of ancestral GFP using a one tailed Mann-Whitney U-test. Each of the five mutations conferred a significantly higher growth rate relative to that of ancestral GFP (Mann-Whitney U-test, false discovery rate corrected *P* < 0.013).

#### Modeling the effect of GroE overexpression on fitness

We define a genotype’s fitness based on its fluorescence rather than its growth rate, because this is the criterion we used during directed evolution. More specifically, we define a genotype’s fitness as the probability that a cell with this genotype exceeds the fluorescence threshold of 150 arbitrary units which we used during phase 1 selection. Cells with a given genotype can show a broad distribution of fluorescence due to phenotypic heterogeneity. Here we show how we map this distribution onto fitness both without and with GroE overexpression. Our procedure consists of two steps. In the first, we estimate statistical parameters such as mean and variance of the fluorescence distribution of each genotype from flow cytometry data. In the second, we use these statistical parameters to predict the fluorescence distribution of genotypes with arbitrary mean fluorescence in the presence and absence of GroE overexpression.

**Step 1**: In the absence of GroE overexpression, all genotypes we engineered showed a Gaussian distribution of logarithmically (base 10) transformed green fluorescence 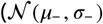, ***Figure S14A-B***). In contrast, in the presence of GroE overexpression this distribution became bimodal. In this bimodal distribution, the first mode (peak) *μ*_+L_ has a lower fluorescence intensity than *μ*_-_ whereas the second mode *μ*_+H_ has a higher fluorescence intensity than *μ*_-_ (***Figure S14A***). The same holds for cyan fluorescence (***Figure S14B***). We expressed this bimodal distribution as a sum of two Gaussian distributions. Specifically, we defined a bimodal probability density function 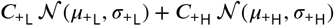, where *C*_+L_ and *C*_+H_ denote weight coefficients for the Gaussian distributions representing the lower and the higher modes, respectively.

To estimate these parameters for each genotype, we fitted a kernel density function to the fluorescence distribution data from populations with GroE overexpression, using the `fitdist` function from the Statistics and Machine Learning Toolbox (ver. 11.5) of MATLAB (2019a). Then, we used the fmincon function (constrained non-linear optimization) from the MATLAB (2019a) Optimization Toolbox (ver 8.3) to estimate a set of parameters for the bimodal distribution that minimize the square distance between the data and the fitted kernel density function. During this optimization, we constrained the weight parameters to have a value between zero and one. In addition, we fitted a (single) Gaussian density function to the fluorescence distribution of GFP mutants in the absence of GroE overexpression, using the `fitdist` function to estimate parameters *μ*_-_ and *σ*_-_ for this distribution.

We performed these calculations for every biological replicate of ancestral GFP and all the engineered mutants, except for start codon mutants. We excluded the start codon mutations from this analysis for two reasons. Firstly, their range of fluorescence intensities (both in the presence and absence of GroE expression) overlapped with the range of cellular autofluorescence, making it difficult to accurately estimate the fluorescence distribution independently from this background. Secondly, for these mutations, the two fluorescence peaks that arose due to GroE expression were so close that bimodality was not clearly apparent. Their overlap with the autofluorescence distribution further hindered the discrimination of these peaks.

Our procedure resulted in an estimate of the parameters *μ*_-_, *σ*_-_, *C*_+L_, *C*_+H_, *σ*_+L_, *σ*_+H_, *μ*_+L_, and *μ*_+H_, for each of the 19 mutants we analyzed, and for three biological replicates for each mutant (***Figure S14***).

Across these mutants, the value of *μ*_+L_ was on average ~ 93% of that of *μ*_-_, and that of *μ*_+H_ was on average ~ 107% of that of *μ*_-_ (***Figure S16A***). In addition, for any one mutation the values of *μ*_+L_ and *μ*_+H_ were clearly distinct from each other (***Figure S16A***). For these reasons, we chose to express *μ*_+L_ and *μ*_+H_ relative to *μ*_-_. By doing that, one can obtain the absolute value of each peak by multiplying the relative values with *μ*_-_. We denote these relative values by the symbols *μ’_+_* _L_ and *μ’_+_* _H_, respectively.

The weight coefficents, *C*_+L_ and *C*_+H_, did not depend on *μ_-_*, and their values showed a nonoverlapping distribution across mutants, with means of 0.64 and 0.76, respectively (***Figure S16B***). The standard deviations *σ_-_*, *σ*_+L_ and *σ*_+H_ also did not depend strongly on *μ_-_* but their distributions across mutants overlapped (***Figure S16C***). Below, we will refer collectively to *C*_+L_, *C*_+H_, *σ*_+L_, *σ*_+H_, *μ*_+L_, and *μ*_+H_ as the parameters of the bimodal distribution.

**Step 2**: To map fluorescence distributions of arbitrary mutants onto fitness, we first represented different mutants through different mean fluorescence values *μ*_-_ in the absence of GroE expression. We explored a range of *μ_-_* values ranging from 10 to 10^5^, because this is the range of green fluorescence that we observed in libraries of GFP mutants before selection. We subdivided this range into 4000 bins that are equally spaced on a logarithmic (base 10) scale, and chose a *μ_-_* value from each bin for all subsequent analyses. (We will henceforth refer to all fluorescence values on this logarithmic scale).

We will first describe how we predicted the fitness of variants for each of these *μ_-_* values in the absence of GroE expression. To do so, we first had to generate a distribution of expected log-fluorescence values for a variant with a given value of *μ_-_*. As mentioned earlier in this section, without GroE expression log-fluorescence is Gaussian distributed with mean *μ*_-_ and standard deviation 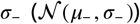. This means that we had to estimate *σ_-_* for any given *μ_-_*. ***Figure S16C*** (black dots) shows that *σ*_-_ spans a range of 0.15 - 0.25 and does not depend on *μ*_-_. We thus chose randomly choose different values of *σ_-_* from this range under the assumption that *σ_-_* itself has a Gaussian distribution. Specifically, we first calculated the mean *(M_σ__*) and standard deviation *(S_σ__*) from the experimentally observed distribution of *σ_-_* values (***Table S3***).

We then chose for each value of *μ_-_*, 4000 different pseudorandom variates *σ_-_* with the Gaussian distribution, 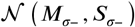. Each of the resulting 4000 pairs of *μ*_-_ and *σ*_-_ values defines a fluorescence distribution of cells with a given genotype, and we determined the fraction of cells in this distribution that exceeded the selection threshold of 150 arbitrary units (2.176 units on a logarithmic scale). We then averaged this fraction over all 4000 *μ_-_-σ_-_* pairs to obtain the expected fitness of a genotype with fluorescence mean *μ*_-_ (F^-^).

We then repeated this procedure for all 4000 values of *μ_-_* in the fluorescence interval (2, 5) to obtain a fitness estimate for each possible variant in this interval. In other words, our estimate of fitness in the absence of GroE is based on 4000×4000 pairs of *μ_-_* and *σ_-_* values.

We next describe how we obtained the same fitness distribution in the presence of GroE expression. We again start with 4000 values of *μ_-_* in the fluorescence interval (2, 5), and perform the same procedure as just described for each such value, except that the distribution is more complex, and we thus need to estimate not just one parameter (*σ*_-_) but six of them: *C*_+L_, *C*_+H_, *σ*_+L_, *σ*_+H_, 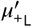 (*μ*_+L_ relative to *μ_-_*), and 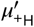 (*μ*_+H_ relative to *μ_-_*). Importantly, our experimental data show these values do not depend on the value of *μ*_-_ (***Figure S16***). We thus estimated each of these parameters by sampling them from a Gaussian distribution whose parameters we estimated from the experimental data (***Table S3***), exactly as we described above for *σ*_-_.

At the end of this procedure we had obtained a total of 4000×4000×6 combinations of parameters. Each of them describes a bivariate fluorescence distribution from which we calculate the fraction of cells above the selection threshold (F^+^).

Overall, this procedure yields for each genotype (value of *μ*_-_) a value of fitness in the absence of GroE (F^-^) and in the presence of GroE (F^+^). We then also calculated the difference between these two fitness values (ΔF = F^+^-F^-^) which denotes the effect of GroE on fitness. A positive value of ΔF means higher fitness in the presence of GroE (buffering) and a negative value means lower fitness in the presence of GroE (potentiation).

## Acknowledgments

This project has received funding from the European Research Council under Grant Agreement No. 739874. We would also like to acknowledge support by Swiss National Science Foundation grant 31003A_172887, by the University Priority Research Program in Evolutionary Biology, as well as by the flow cytometry facility and the functional genomics center at the University of Zurich. We thank Andrei Papkou for his suggestions on statistical analyses, and Miriam Olombrada Sacristan, Jia Zheng and Shraddha Karve for their assistance during the experiments.

## Data files

All data are available in the manuscript or supplementary materials. SMRT sequencing data are available at NCBI Sequence Read Archive (SRA) under the BioProject ID, PRJNA706377.

## Supplementary Online Material

## 1 Construction of the expression system

**Figure S1:**
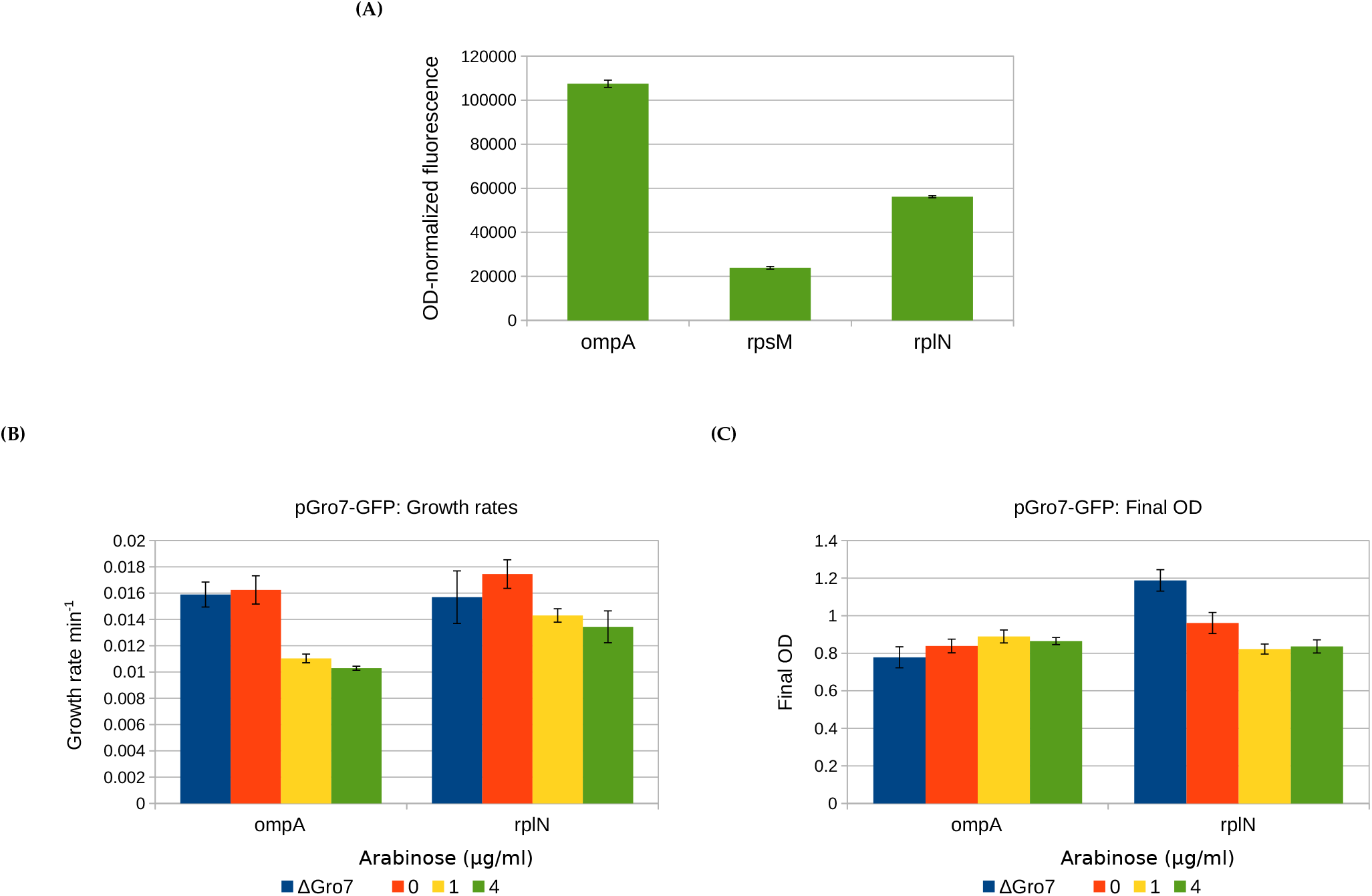
Effect of different promoters on fluorescence and growth. **(A)** Steady state fluorescence of pGro7 strains with GFP expressed from the ompA, rpsM and rplN promoters. Growth rate **(B)** and final OD **(C)** of pGro7 strains with induction at different concentrations of arabinose. Δ denotes the corresponding pΔGro7 strain.

**Figure S2:**
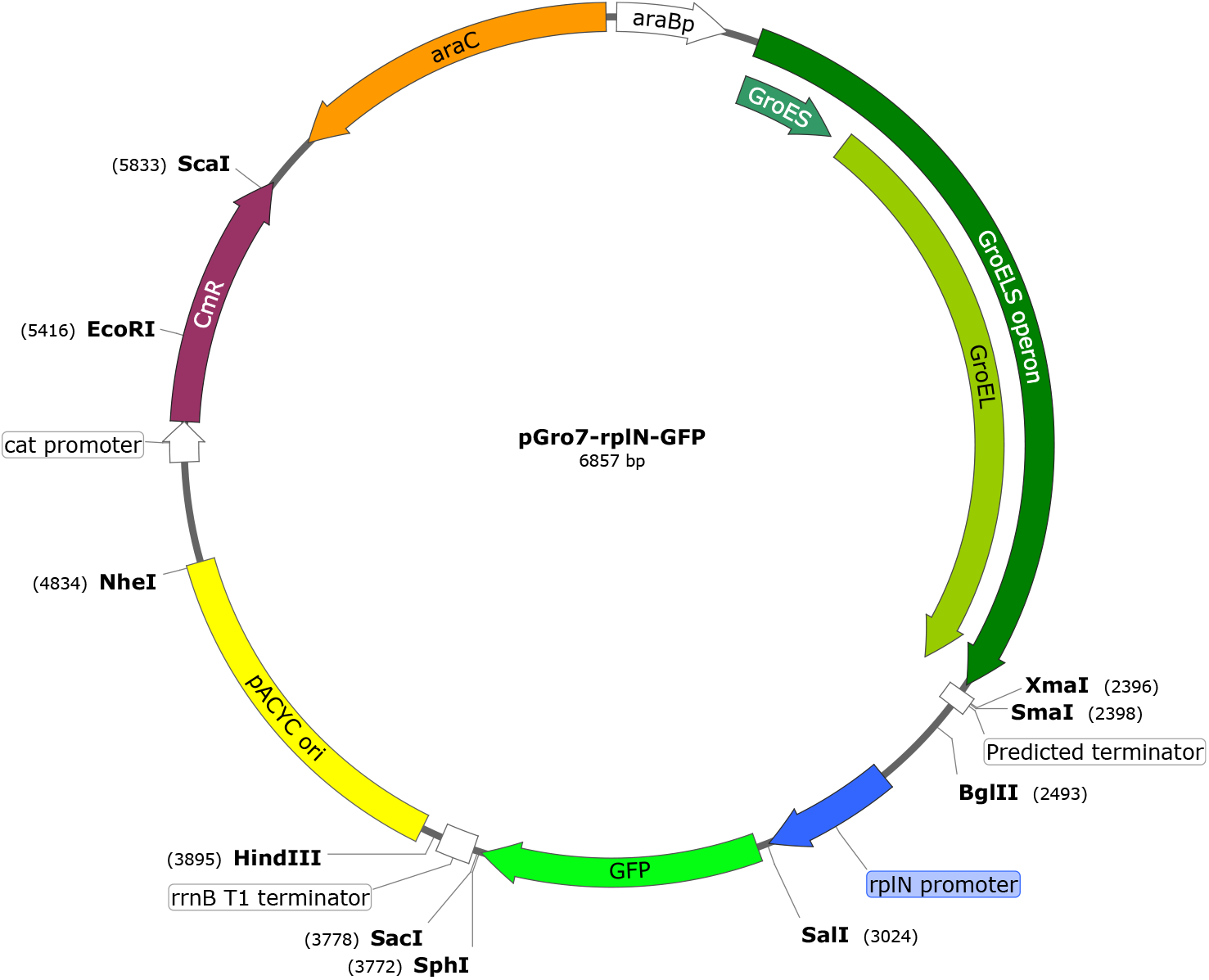
Plasmid map of pGro7-rplN-GFP.

**Figure S3:**
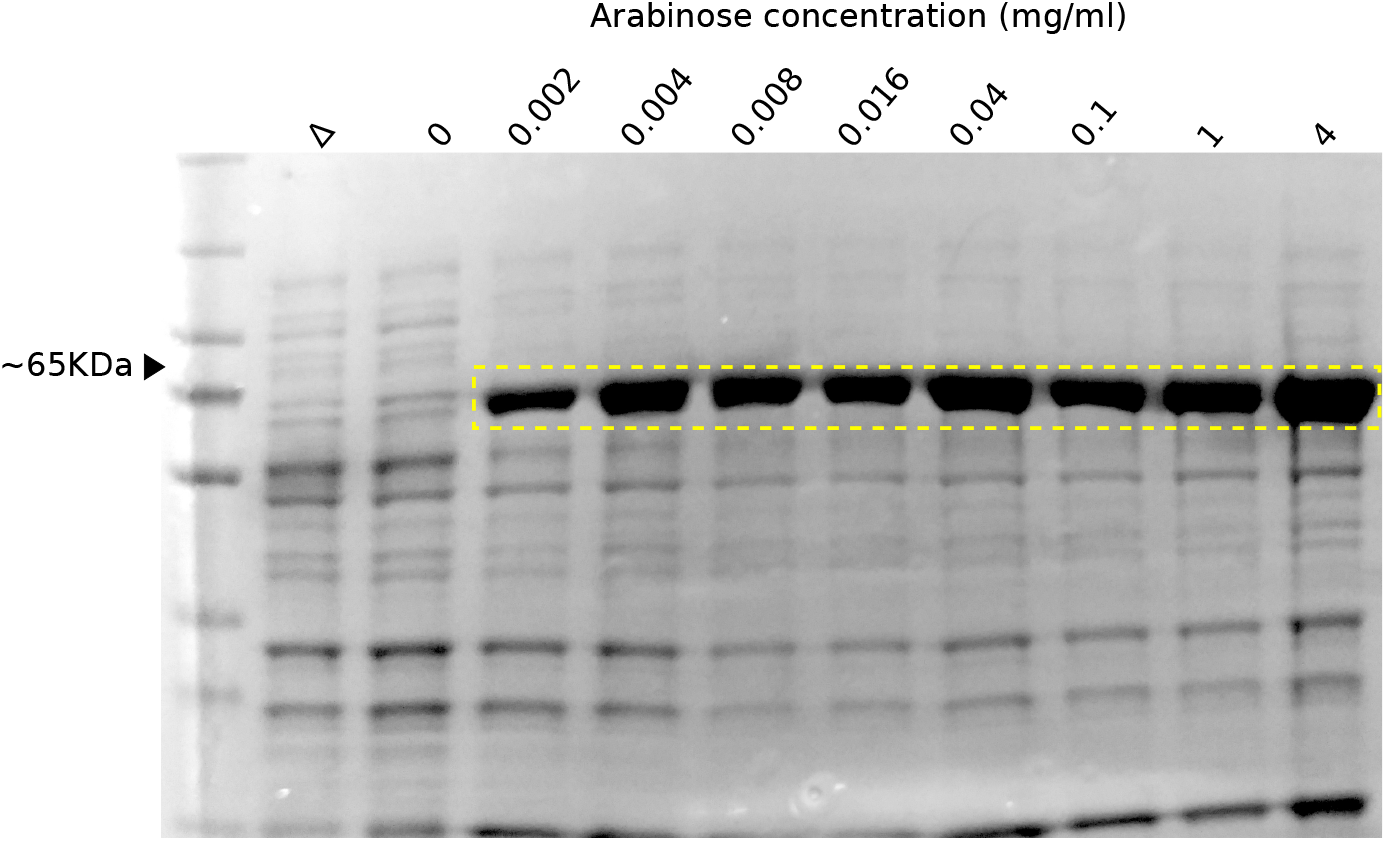
Image of polyacrylamide gels post-electrophoresis, showing the protein profile of cells with GroE expression induced at different concentrations of arabinose (on top of the gel). The symbol Δ on top of the second lane from the left denotes protein extracts from the pΔGro7-rplN-GFP strain. Protein molecular weight markers may not denote the corresponding weights accurately. The region boxed in yellow contains bands corresponding to GroEL (~60KDa).

## 2 Ancestral GFP is a GroE client

In this analysis, we determined whether GroE expression affected the fluorescence of the ancestral GFP. To this end, we measured the fluorescence of 10,000 cells expressing the ancestral GFP, with and without GroE expression, using flow cytometry (Methods). We found that GroE expression created phenotypic heterogeneity in GFP expression, which is evident by a bimodal distribution of fluorescence (Figure S4). This change in the fluorescence distribution suggests that our GFP variant is likely to be a GroE client. One of the two fluorescence intensity peaks has a higher intensity and the other one has a lower intensity than the unimodal peak of the fluorescence distribution without GroE induction.

**Figure S4:**
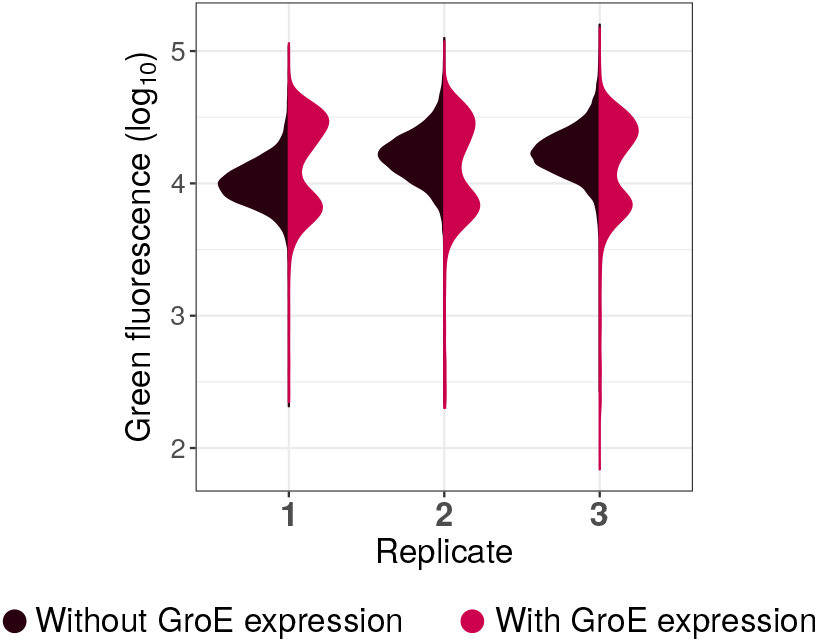
GroE expression affects the fluorescence of ancestral GFP. Violin plots denote the distribution of green fluorescence (log_10_ transformed; vertical axis) of the ancestral GFP protein with (red) and without (blue) GroE expression. We performed these measurements in three biological replicates shown in the horizontal axis.

## 3 Populations evolve towards a new phenotype (cyan fluorescence) during phase 2

**Figure S5:**
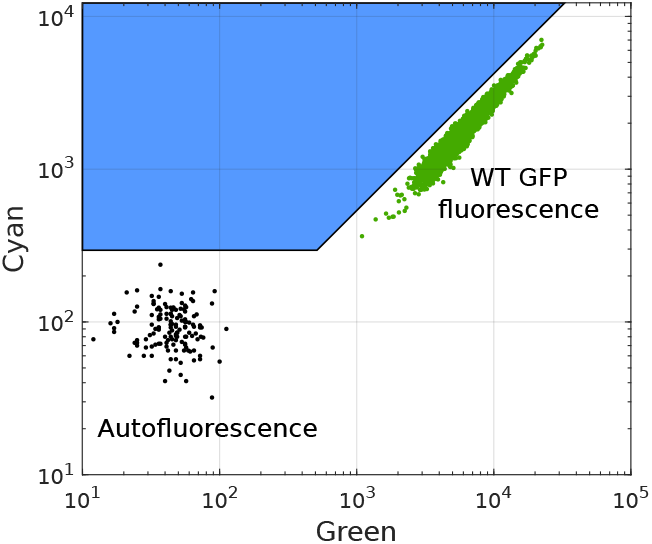
Strategy for selection of cells exhibiting a shift towards cyan fluorescence. Green circles denote the fluorescence of cells expressing wild type GFP, and black circles denote the autofluorescence of cells. We selected cells mapping to the blue area during directed evolution in phase 2.

**Figure S6:**
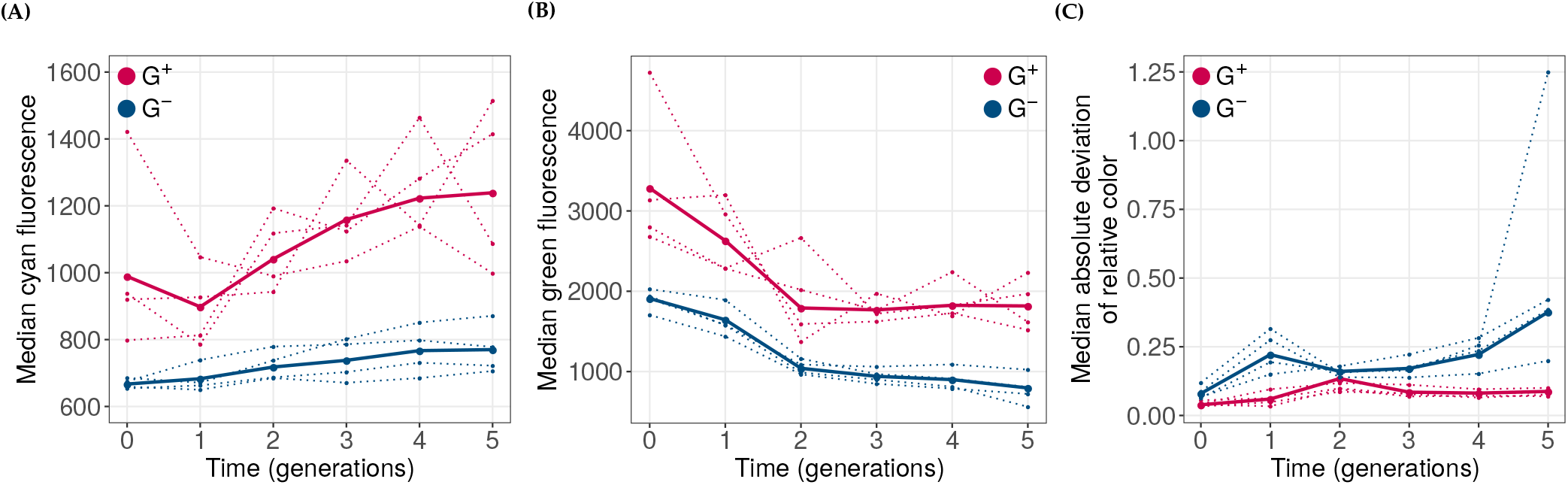
Effect of the GroE overexpression on the evolution of the novel phenotype. **(A)** Cyan and **(B)** green fluorescence (arbitrary units, vertical axes) of evolving G^+^(red) and G^-^ (blue) populations during phase 2 of directed evolution. **(C)** The median absolute deviation of relative color (ratio of cyan and green fluorescence, vertical axis) in the evolving populations. In all panels, dotted lines denote the median values of individual replicate populations, and solid lines denote the median values when data from all populations is pooled. The horizontal axis denotes the time in generations (round of evolution) where generation zero refers to populations at the end of phase 1 evolution.

## 4 GroE may buffer a few mutations in phase 2 populations

The loss of genetic diversity (Main text: Figure 8) in phase 2 populations suggests that GroE potentiates the effect of deleterious mutations overall. However, this does not exclude the possibility that it buffers the effect of some such mutations. Such mutations should preferentially persist in G^+^ populations and their effect on fluorescence should increase with GroE expression. Specifically, if buffering takes place, G^+^ populations at the end of phase 2 should increase their fluorescence when GroE is overexpressed compared to when it is not overexpressed. This is indeed the case (Figure S7). Cyan fluorescence was significantly higher in these populations when we overexpressed GroE (Mann-Whitney U-test, *P* < 6.5 × 10^-4^), although the extent of this increase was small (2 - 25%; Figure S7).

This suggests that at least some buffering of mutations takes place during phase 2.

**Figure S7:**
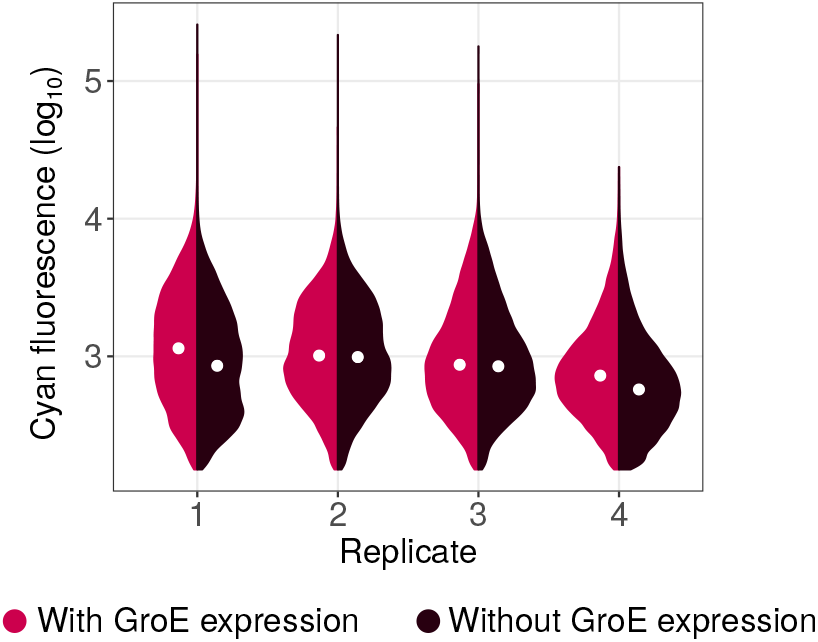
Violin plots denoting the distribution of logarithmically (base 10) transformed cyan fluorescence (arbitrary units) for each replicate of G^+^ populations at the end of phase 2 evolution (generation 5), with (red) or without (brown) the expression of GroE. The white circle in the center of the distribution denotes the median. The medians are significantly different for each pair of distributions shown (Mann-Whitney U-test, *P* < 6.5 × 10^-4^).

## 5 GroE expression affects the spectrum of accumulated genotypes in both phase 1 and phase 2

To find out whether the chaperone causes different kinds of genotypes to accumulate in phase 1 populations, we randomly sampled 200 sequences from each population at the end of phase 1, and displayed the location of these sequences in genotype space using PCA (Materials and Methods: Principal component analysis of the genotypes). Populations that evolved under chaperone overexpression cluster in different regions of genotype space than the control populations (Figure S8A). These patterns are corroborated by a complementary PCA, which we conducted on the frequency spectrum of alleles that differ in specific amino acids from the ancestor (Figure S8B).

We performed a similar analysis for populations at the end of phase 2 and found that G^+^ populations accumulate a different set of genotypes compared to G^-^ populations (Figure S9).

**Figure S8:**
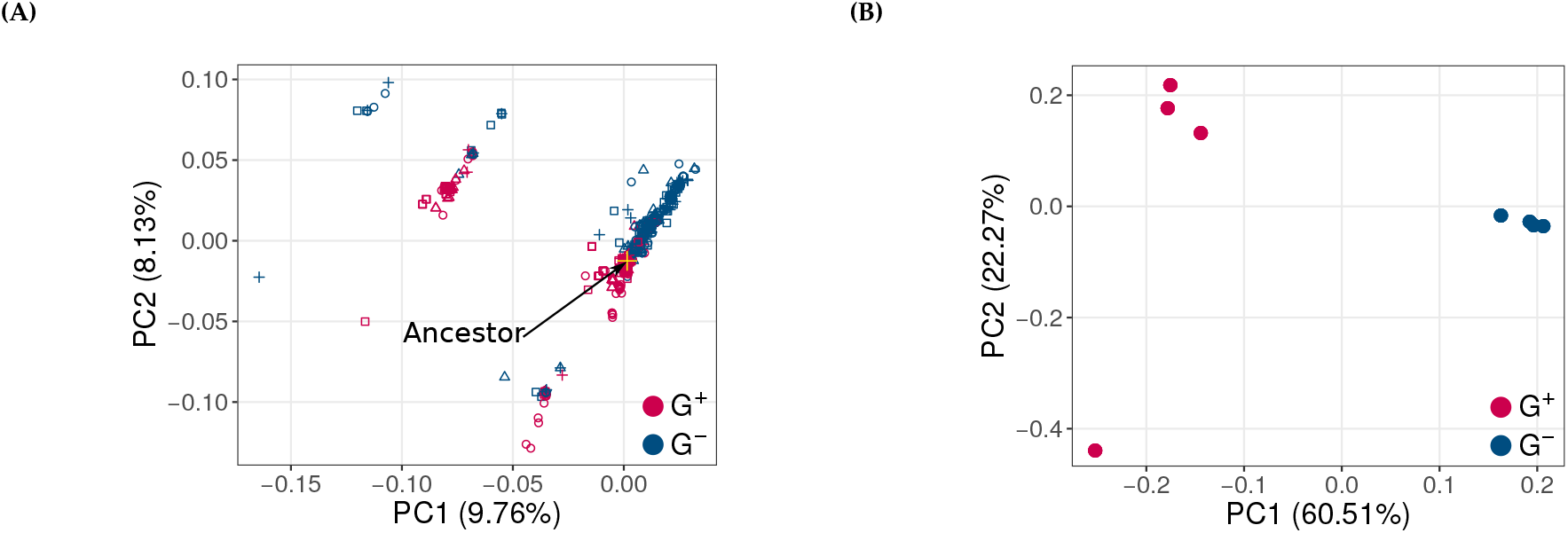
Effect of GroE on the genotype of populations at the end of phase 1 evolution. **(A)** Principal component analysis (PCA) of genotypes in G^+^ (red) and G^-^ (blue) populations. The ancestral genotype is represented by a yellow cross. **(B)** PCA based on the frequencies of individual mutations in G^+^ populations (red) and G^-^ (blue) populations.

**Figure S9:**
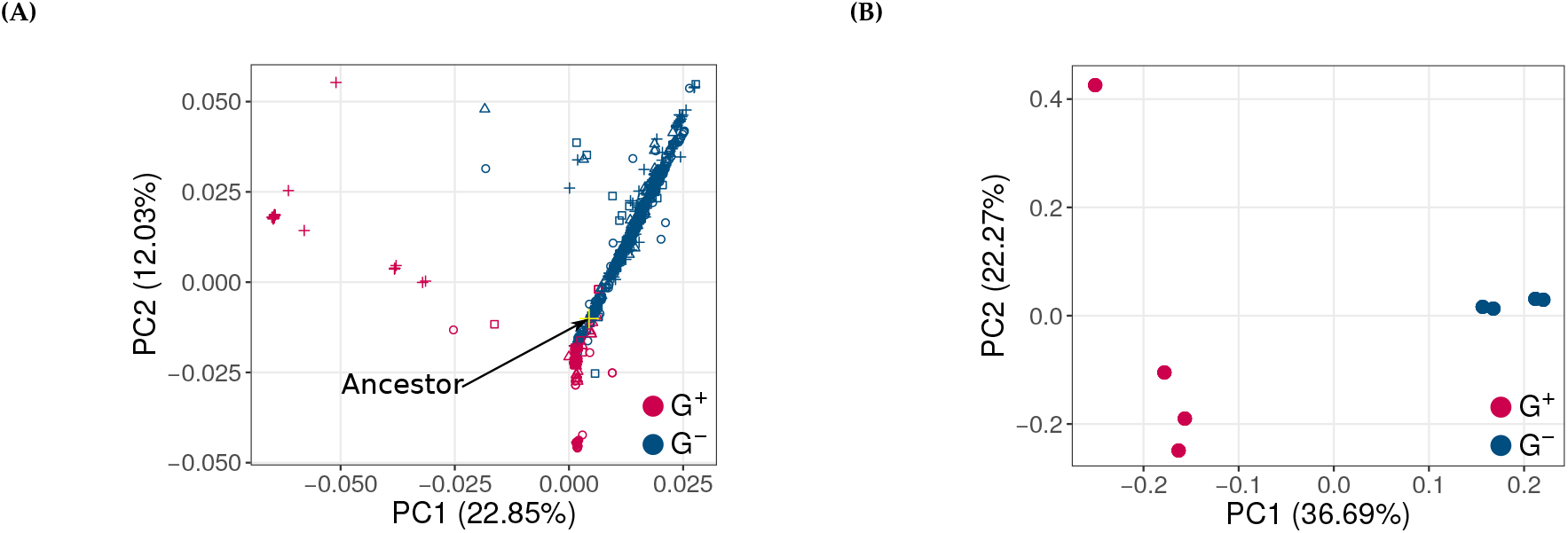
Effect of GroE on the genotype of populations at the end of phase 2 evolution. **(A)** Principal component analysis (PCA) of genotypes from G^+^ (red) and G^-^ (blue) populations. The ancestral genotype is represented by a yellow cross. **(B)** PCA based on the frequencies of individual mutations in G^+^ populations (red) and G^-^ (blue) populations.

## 6 GroE affects the accumulation of single amino acid mutations

**Figure S10:**
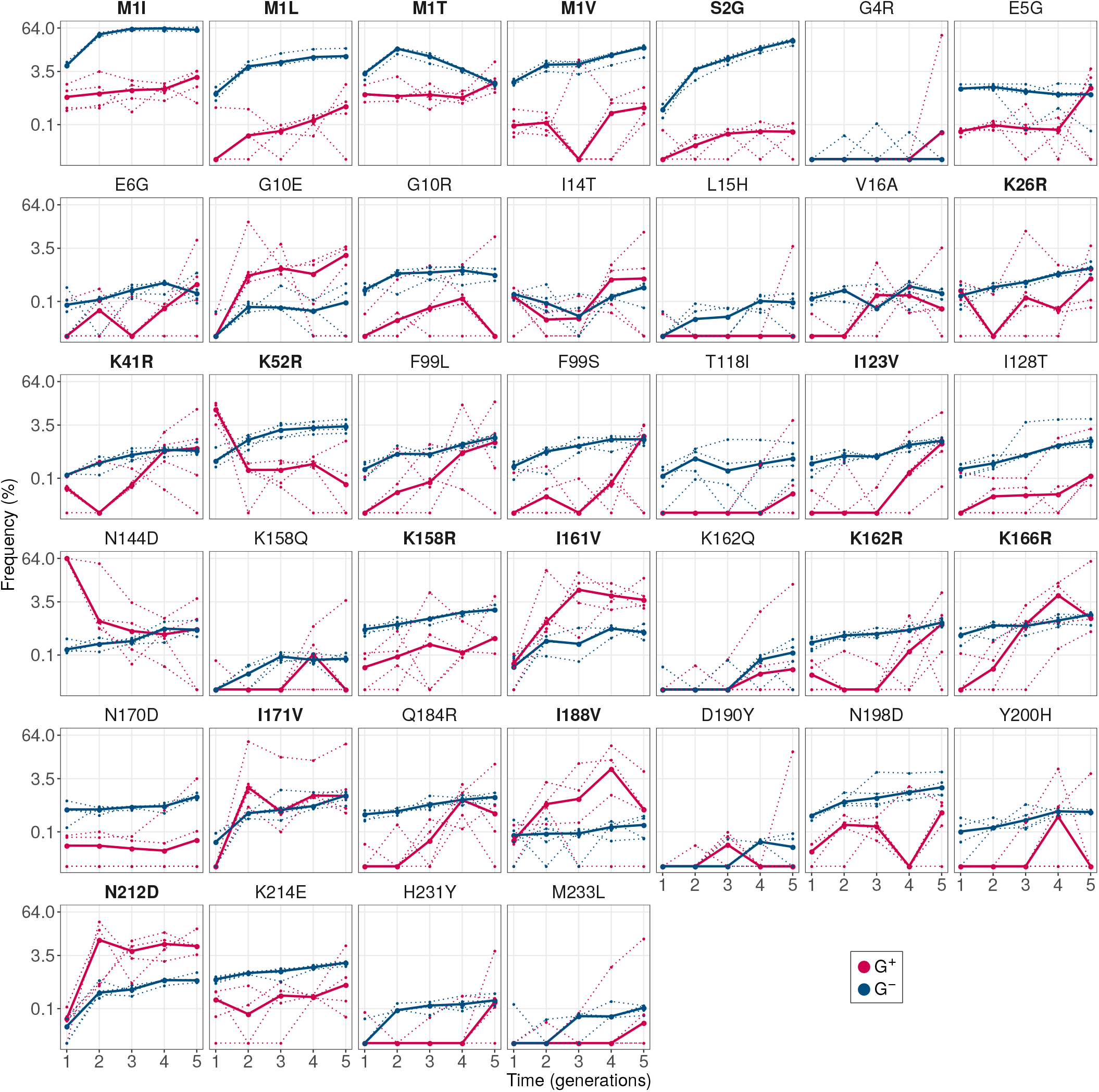
We show only those mutations whose frequencies exceed 3.5% in at least one replicate population by the end of evolution, and are differentially enriched between G^+^ (red) and G^-^ (blue) populations. Mutations that we analyzed further are highlighted with bold mutant names. Vertical axes (in log scale) denote the frequency whereas the horizontal axes denote the generation i.e. round of evolution. Dotted lines denote the frequency of mutations in individual replicates whereas solid lines denote the median frequency.

**Figure S11:**
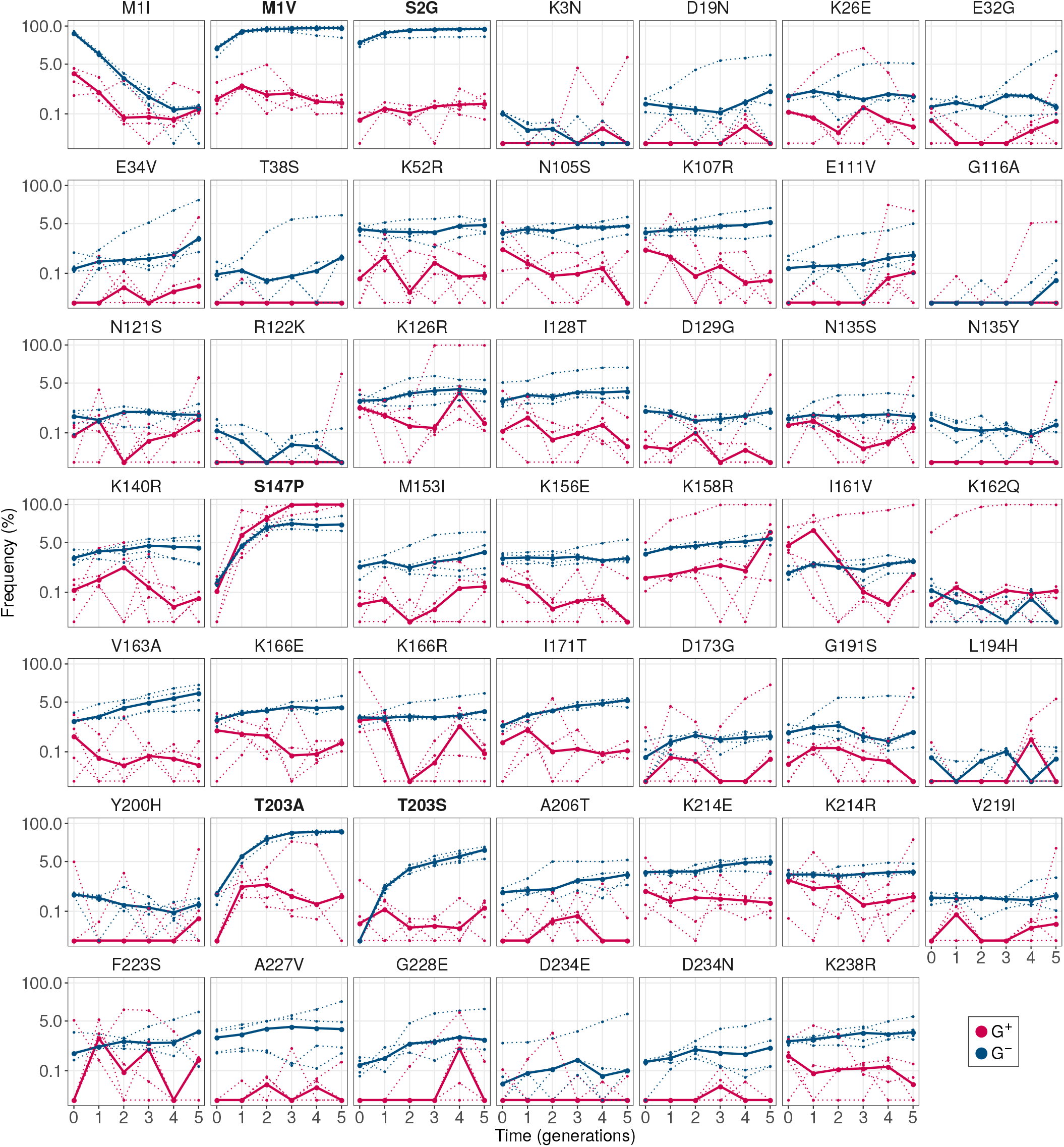
Frequency of single amino acid mutations during phase 2 evolution. We have shown only those mutations whose frequencies exceed of 5% in at least one replicate population, by the end of evolution and are differentially enriched between G^+^ (red) and G^-^ (blue) populations. We included M1I in these plots to show that while it is abundant in phase 1 G^-^ populations, it is lost in phase 2. Mutations that we analyzed further are highlighted with bold face font. Vertical axes (in log scale) denote the mutation frequency and the horizontal axes denote the generation i.e. round of evolution. Dotted lines indicate the frequency of mutations in individual replicates, whereas solid lines denote the median frequency.

## 7 Deleterious mutations do not hitchhike with potentially stabilizing mutations in the absence of GroE expression.

Most random mutations are likely to be deleterious^[1–3]^ and they can persist in populations by hitchhiking with beneficial or stabilizing mutations. The high frequency of the deleterious mutations M1I, M1L, M1V and S2G in G^-^ populations raises the possibility that these mutations hitchhike with other beneficial mutations. To exclude this possibility, we analyzed the frequencies of variant genotypes that harbor one or more single mutation. We found that the most abundant genotype in all the G^-^ populations contained just the mutation M1I (Table S1). This genotype exceeded a frequency of 10% in all the replicate populations. We next determined if the other three abundant deleterious mutations (M1L, M1V and S2G) also frequently existed as single mutation genotypes. This was indeed the case. Specifically, these single mutation genotypes were among the six most frequent genotypes in all replicate populations, and had a frequency that ranged between 0.45 and 40 percent (Table S1). This suggests that these deleterious mutations can rise to appreciable frequencies on their own. We next analyzed the multi-mutation genotypes in G^-^ populations and found that M1I+S2G and M1V+S2G were the most abundant multi-mutation genotypes in every replicate population, with a frequency range of 1.5 – 5%. No other multi-mutation genotype rose above the frequency of 0.9% in any replicate population. To further validate these findings, we calculated the pairwise cooccurrence frequency of different mutations in G^-^ populations. Consistent with our previous finding, S2G coexisted most frequently with M1I and M1V in more than 6.5% of the genotypes and in every replicate population. In contrast, no other mutation pair had a frequency higher than 4% in any replicate population. Taken together, these results indicate that the abundance of deleterious mutations in G^-^ populations is not due to hitchhiking.

**Table S1:** Frequencies of the 10 most frequent genotypes in G populations at the end of phase 1.

| Replicate 1 |  | Replicate 2 |  | Replicate 3 |  | Replicate 4 |  |
| --- | --- | --- | --- | --- | --- | --- | --- |
| Genotype | Frequency | Genotype | Frequency | Genotype | Frequency | Genotype | Frequency |
| M1I | 12.13 | M1I | 10.13 | M1I | 16.39 | M1I | 10.91 |
| M1V, S2G | 4.89 | M1L | 3.98 | M1I, S2G | 2.82 | M1V, S2G | 2.97 |
| M1I, S2G | 2.77 | M1V, S2G | 2.97 | M1L | 2.42 | M1L | 2.09 |
| M1L | 1.86 | M1I, S2G | 2.18 | M1V, S2G | 1.46 | M1I, S2G | 2.09 |
| M1V | 1.12 | S2G | 1.12 | S2G | 1.01 | M1V | 1.28 |
| S2G | 1.06 | M1V | 1.06 | M1I, I167T | 0.84 | S2G | 0.88 |
| M1I, V163A | 0.53 | M1I, N198D | 0.9 | M1I, K52R | 0.51 | M1I, I128T | 0.8 |
| M1T | 0.43 | M1T | 0.5 | M1V | 0.45 | M1T | 0.4 |
| M1I, K214E | 0.43 | M1I, N105S | 0.5 | M1I, G191A | 0.45 | M1I, N198D | 0.4 |
| M1I, K79R | 0.37 | M1L, N198D | 0.39 | M1I, N105S | 0.45 | M1I, N170D | 0.4 |

**Table S2:** Frequencies of the 10 most frequent genotypes in G^+^ populations at the end of phase 1.

| Replicate 1 |  | Replicate 2 |  | Replicate 3 |  | Replicate 4 |  |
| --- | --- | --- | --- | --- | --- | --- | --- |
| Genotype | Frequency | Genotype | Frequency | Genotype | Frequency | Genotype | Frequency |
| F99L | 5.37 | G10R | 2.32 | G4R | 31 | K166R, I171V | 27.19 |
| I161V, N212D | 4.21 | K162Q, M233L | 1.92 | K158R | 1.38 | D190Y | 14.29 |
| I14T, F99L | 2.95 | K101E | 1.92 | N212D | 1.09 | K166R | 10.65 |
| E6G | 2.74 | K162Q | 1.51 | M1T | 1.09 | K166R, D190Y | 1.82 |
| N212D | 1.84 | M1T | 1.21 | K158R, N212D | 1.09 | K166R, I171S | 0.91 |
| N105S | 1.74 | K41R | 1.11 | F99S | 0.86 | I161V, N212D | 0.81 |
| K26R | 1.69 | Y200H | 1.11 | V120I, N144D | 0.8 | E115G | 0.81 |
| I14T | 1.69 | N170D, I188V | 0.91 | F99S, V163A | 0.75 | I171V | 0.76 |
| I123V, I161V, Q184R, N212D | 1.37 | I188V | 0.81 | P211T | 0.75 | N212D | 0.67 |
| T38A | 1.16 | E5G | 0.71 | I123V | 0.75 | K79R, I161V, N212D | 0.62 |

## 8 Deleterious mutations rarely accumulate through GroE mediated buffering

Because the phenotype of populations evolved under GroE overexpression (G^+^) is not strongly dependent on chaperone expression (Main text: Figure 4), the presence of most deleterious mutants in these populations cannot be explained by GroE mediated buffering. To further validate this observation, we analyzed mutations that are enriched in G^+^ populations, i.e. mutations that have a higher frequency in these populations compared to G^-^ populations. They are the most promising candidates for mutations whose deleterious effects are buffered by GroE. In this analysis, we focused on mutations that attained a frequency exceeding 3.5% in at least one replicate population. We found 25 such mutations with significantly higher frequency in G^+^ populations (GLM: likelihood ratio test [LRT], P < 10^-3^) in G^+^ populations after the end of phase 1 evolution (Figure S10). The majority of these 25 mutations met our selection criterion in only one out of four populations. Moreover, in some replicate populations these mutations initially increased in frequency only to become lost again (frequency less than 0.01%). With the exception of I161V and N212D (Figure S10) the mutations did not steadily increase in frequency. Thus, most mutations enriched in G^+^ populations did not show the kind of evolutionary dynamics expected from consistent and sustained buffering.

We nonetheless analyzed selected mutations in more detail to study their phenotypic effects. They include two classes of G^+^-enriched mutations that stand out because of a common characteristic. These are isoleucine to valine (I→V) and lysine to arginine (K→R) mutations. We identified four G^+^-enriched mutations in each class. These are I123V, I161V, I171V, I188V, K26R, K41R, K162R and K166R. In addition, one start codon mutation (M1T) was also G^+^- enriched. This mutation provides a contrast to our previously analyzed start codon mutations that were deleterious, potentiated, and thus G^-^-enriched (Main text: Figure 5). We hypothesized that M1T was deleterious and possibly buffered. Furthermore, we also analyzed the mutation N212D, because it was one of the only two mutations that steadily increased in frequency during phase 1 evolution. (The other is I161V and it is included in the list of isoleucine to valine mutations we analyzed.)

To identify how each of these 10 mutations affected the phenotype, we engineered them individually into ancestral GFP, and quantified their fluorescence in three biological replicate measurements. Since N212D was one of the only two mutations that appeared to have steadily increased in frequency during phase 1 evolution, we also engineered this mutation and quantified its fluorescence. We found that M1T was indeed deleterious for fluorescence, reducing it by 65 - 69 fold relative to ancestral GFP. Of the other nine mutations, only two (K166R and I171V) showed a modest (7 - 40%) increase in fluorescence relative to ancestral GFP in all the three biological replicate measurements we conducted. The remaining seven mutations did did not significantly affect fluorescence (Figure S12).

**Figure S12:**
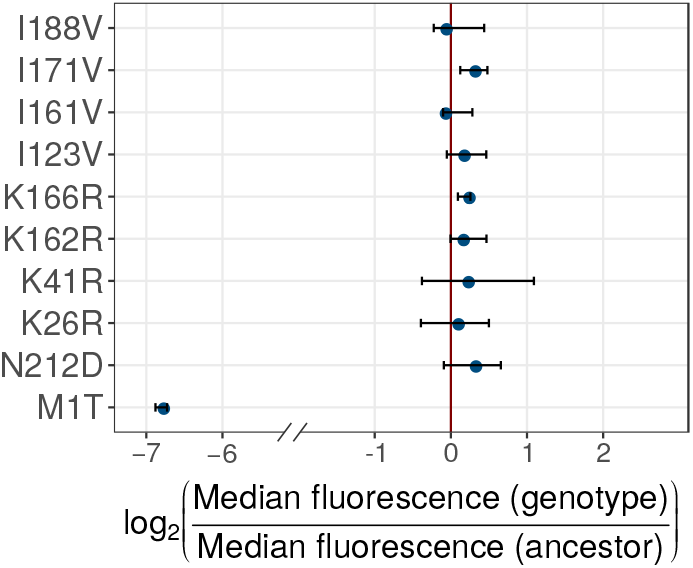
Fluorescence of variants enriched in G+ populations at the end of phase 1. The figure shows the fluorescence of mutations that are more frequent in G^+^ populations than in G^-^ populations. The horizontal axis shows the log_2_-transformed ratio between the median population green fluorescence for a given GFP variant (vertical axis) and the median green fluorescence of ancestral GFP. A negative value denotes a deleterious effect whereas a positive value denotes a beneficial effect. The solid blue circle denotes the median value of the log_2_ transformed fluorescence ratios in the three replicate measurements whereas the errorbar spans the range of minimum and maximum values.

## 9 GroE delays the accumulation of color shifting mutations in phase 2 populations

During phase 2 GroE delays the evolutionary change in fluorescence color shift (Main text: Figure 7, Figure S6C). To find out which mutations may be involved in this change, we first identified single amino acid variants in the evolving populations that had very low frequency in phase 1 but high frequency in all replicate populations of phase 2, reasoning that these variants may be involved in the color shift. (Figure S11). The three variants that satisfied this criterion are S147P, T203A and T203S (Figure S13). We engineered these variants into ancestral GFP to study their fluorescence. In addition, we engineered the double variants S147P/T203A and S147P/T203S. All three single variants and both double variants shifted color, i.e., they showed a significantly higher cyan fluorescence relative to green fluorescence than ancestral GFP (Mann-Whitney U-test, *P* < 10^-15^, Figure S15A). Of the single variants, T203S showed the greatest color shift (1150% of the ancestral relative color) followed by S147P (130%) and T203A (105%). Moreover, the two single amino acid changes in the double variants had a synergistic effect, such that the relative color of the double variant was higher than the sum of the individual single variant (Figure S15B).

**Figure S13:**
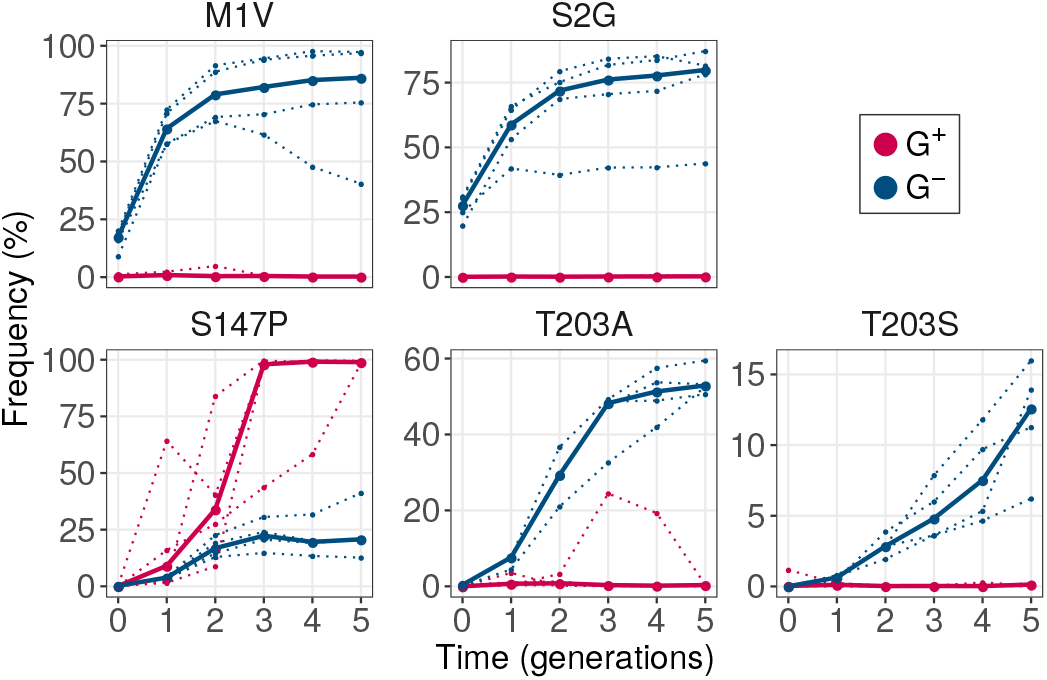
GroE overexpression disfavors accumulation of deleterious mutations and several color-shifting mutations in phase 2. The vertical axes show mutation frequency as a function of time (generations, horizontal axes). Each panel shows the evolutionary dynamics of a different mutant in evolving G^+^ (red) and G^-^ (blue) populations. M1V and S2G are deleterious. S147P, T203A and T203S are color-shifting and beneficial. In each panel, dotted lines denote mutation frequencies in individual replicate populations, and the solid line denotes the median frequency.

Importantly, while G^-^ populations accumulated all three single mutations and the double mutations at varying frequencies, G^+^ populations only accumulated the variant S147P, which swept through these populations. This can explain why the color shift is less pronounced in G^+^ populations. To explain how GroE expression could cause this phenomenon, we considered three hypotheses. First, the color-shifting variants may be deleterious and hence disfavored by GroE (Main text: Figure 6). This is not the case. Both T203A and T203S (also S147P) and both double mutants are in fact beneficial, i.e., they increase the absolute cyan fluorescence (Figure S15C). Second, GroE might reduce the fluorescence of the T203A/S mutants and thus disfavor their selection. To find out whether this is the case, we measured the fluorescence of all three single mutants and the two double mutations under GroE expression. We found that GroE expression led to the kind of phenotypic heterogeneity we previously discussed for phase 1 mutations (Figure S14B). Like phase 1 mutations, the distribution of log transformed fluorescence, which was uni-modal without GroE expression, became bimodal upon GroE expression. One of the two bimodal peaks had lower fluorescence intensity (*μ_+L_*) than the unimodal peak (*μ_-_*) whereas the other peak (*μ_+H_*) had a higher fluorescence intensity than μ-. As we noted previously using our computational model, GroE expression does not affect the fitness of non-deleterious mutations, thus ruling out the possibility that GroE reduces the fluorescence and thereby the fitness of T203A/S. Third, GroE expression might itself reduce the color shift in T203A/S and thereby disfavor the accumulation of these variants in G^+^ populations. To find out, we compared the relative color of each genotype under GroE expression with that of the ancestral GFP (without GroE expression. We found that the relative color of T203A, T203S, S147P, and both double mutants was still significantly higher under GroE expression than that of ancestral GFP in the absence of GroE expression(Mann-Whitney U-test, P < 10^-15^). This rules out the possibility that GroE expression suppresses the color shift of these mutations.

In sum, while it is clear that GroE can reduce the fluorescence color shift during evolution by affecting the spreading of color-shifting mutations, the reasons for this observation remain a task for future work.

## 10 Fluorescence analysis of engineered mutants

We measured the fluorescence of each GFP mutants with and without GroE expression, using flow cytometry (Materials and Methods: Analysis of fluorescence of populations using flow cytometry). For each mutant, we measured the fluorescence of 10,000 cells to obtain a fluorescence distribution. Also for each mutant, we performed these measurements in three biological replicates, repeating the measurement starting from three different samples of the same mutant’s glycerol stock. We also re-measured fluorescence of ancestral GFP along with that of the mutants for every biological replicate experiment, to correct for day to day variations in growth conditions and performance of the flow cytometer. We note that GroE expression led to heterogeneity in fluorescence for most mutants and replicates (Figure S14A-B), as we had observed for ancestral GFP (Figure S4).

To analyze the spectral shift in fluorescence in any one genotype harboring one or more mutations, we first calculated the ratio of cyan fluorescence and green fluorescence (relative color) for every cell in an isogenic population of this genotype, in the absence of GroE expression. In this way, we obtained the distribution of relative color for every variant genotype. Next, we compared the distribution of relative color of every genotype with that of the ancestral GFP using Mann Whitney U-test. We found that the genotypes S147P, T203A, T203S, S147P/T203A and S147P/T203S showed a significant increase in relative color (Mann Whitney U-test, P < 10^-15^; Figure S15).

Next, we asked if GroE expression reduces the color shift of some variants, thereby disfavoring their selection in phase 2 evolution. To this end, we compared the relative color of each genotype under GroE expression with that of the ancestral GFP (without GroE expression), using a Mann-Whitney U-test as described in the previous paragraph. This analysis showed that the relative color of each mutant genotype was still significantly higher than that of ancestral GFP (Mann Whitney U-test, P < 10^-15^).

**Figure S14:**
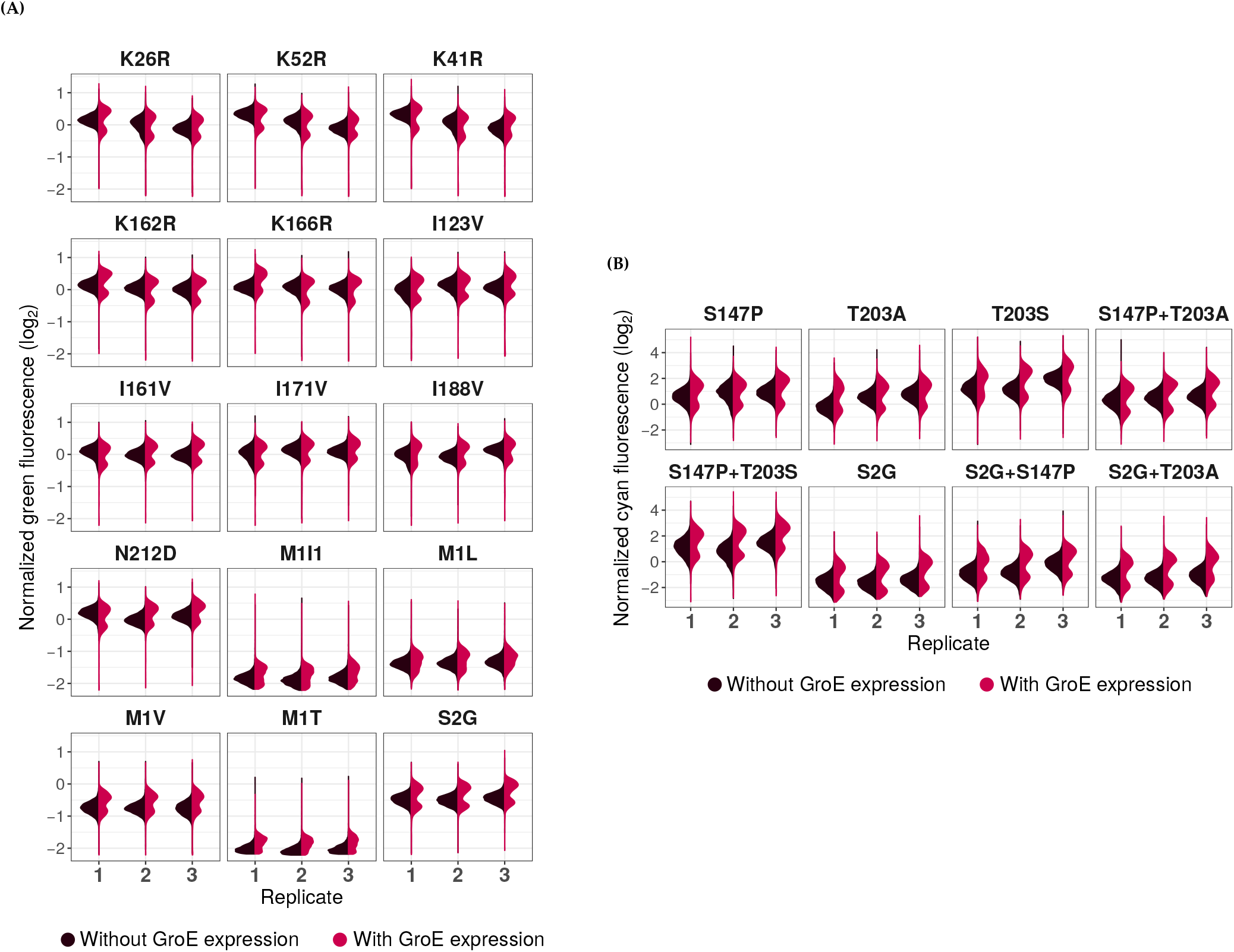
GroE overexpression causes phenotypic heterogeneity in populations expressing different GFP mutant genotypes. For each biological replicate, we normalized the fluorescence of a given mutant genotype to the median fluorescence of ancestral GFP measured at the same time. This also allows us to visualize the fluorescence distribution of a genotype with respect to the ancestral GFP. The violin plots show the distribution of normalized fluorescence of genotypes in the presence (red) and the absence (blue) of GroE expression. Panel **(A)** shows the distribution of normalized green fluorescence of selected differentially enriched mutations after phase 1 evolution, and panel **(B)** shows the distribution of normalized cyan fluorescence of selected differentially enriched mutations after phase 2 evolution.

**Figure S15:**
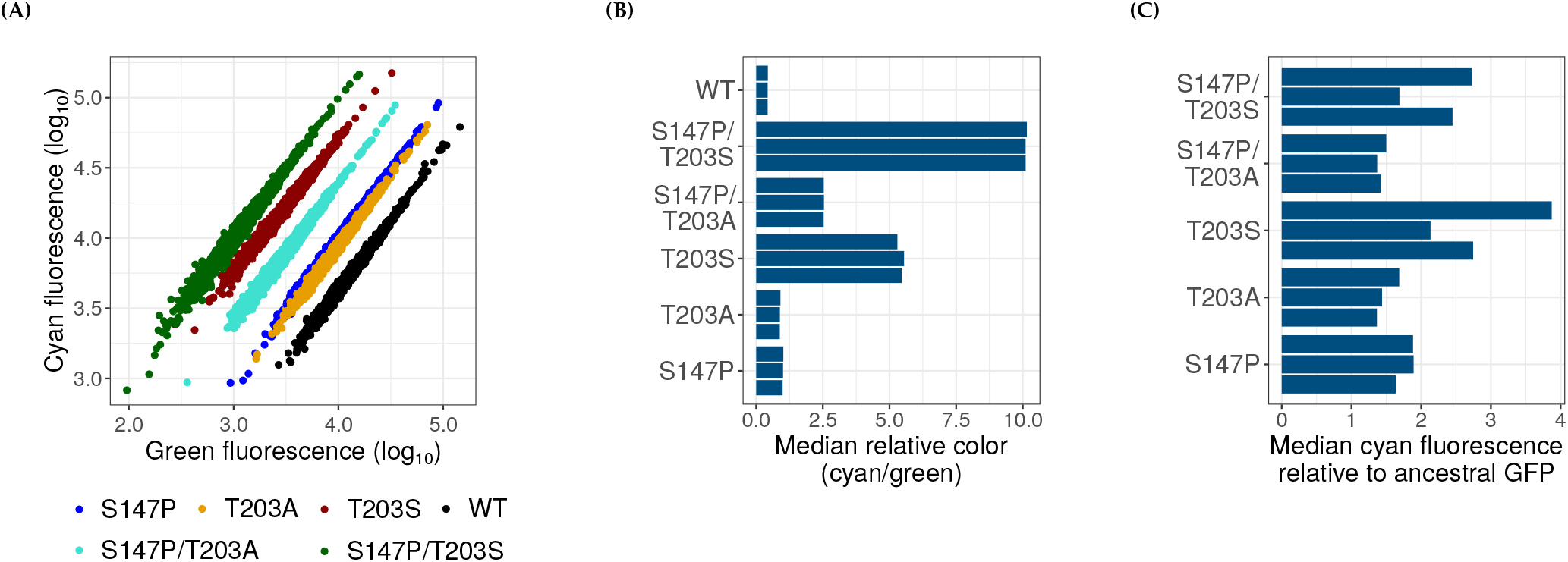
Beneficial color shifting mutations accumulate in phase 2. **A** Single mutants, S147P (blue), T203A (yellow) and T203S (brown), as well as the double mutants S147P/T203A (turquoise) and S147P/T203S (dark green) show a color shift towards cyan fluorescence. Ancestral GFP (black) is denoted by the symbol WT. The horizontal axis shows log_10_ transformed green fluorescence and the vertical axis shows log_10_ transformed cyan fluorescence. **(B)** Median relative color (cyan:green ratio; horizontal axis) for different variants (vertical axis) is in agreement with the scatterplot. **(C)** All the mutations (vertical axis) i.e. the three single mutations and the two double mutations are beneficial as they increase the median cyan fluorescence (horizontal axis). Three grouped bars in panels **B-C** corresponding to each mutation denote three biological replicate measurements.

## 11 GroE expression affects the fitness of different variants

We estimated the fitness of different variants represented by a unique value of μ- (mean of Gaussian distributed log_10_-transformed fluorescence in the absence of GroE expression; Materials and Methods: Modeling the effect of GroE overexpression on fitness). The average fitness in the presence and absence of GroE expression (F^+^ and F^-^, respectively) increased with increasing μ- (Figure S17A). However, also with increasing values of μ-, the effect of GroE (ΔF) increased from zero, changed its sign from positive to negative, and eventually increased again to approach zero from below. This indicates that GroE buffers some mutations, potentiates other mutations, and has no effect on the fitness of the rest (Main text: Figure 6).

**Figure S16:**
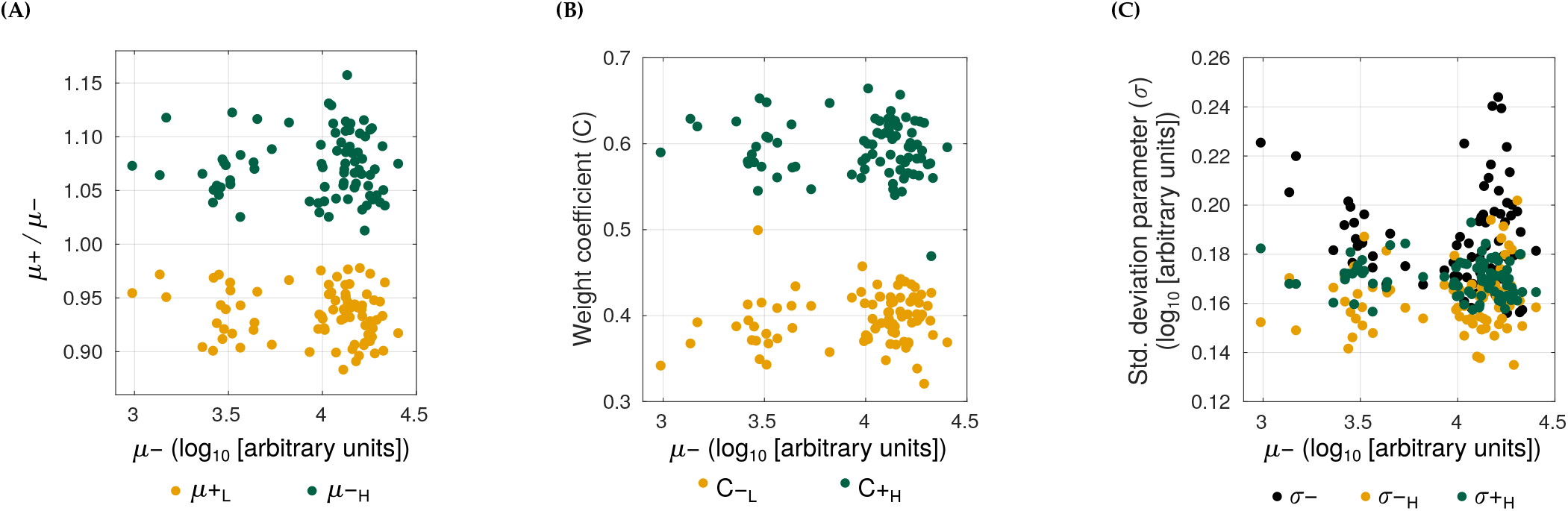
Similar qualitative changes in fluorescence distribution for different mutations due to GroE expression. Each panel shows a scatterplot of the correlation between *μ_-_* (horizontal axes) and other parameters that characterize the fluorescence distribution (vertical axes). In panel **(A)** the vertical axis shows the position of the bimodal fluorescence intensity peaks μ_+L_ (yellow) and μ_+H_ (green) relative to μ-. In panel **(B)** the vertical axis shows the weight coefficients C_+L_ (yellow) and C_+H_ (green), and in panel **(C)** the vertical axis shows the peak width parameters *σ_-_* (black), σ_+L_ (yellow), and σ_+H_ (green). In all three plots, each data item (circle) is derived from the distribution of log_10_-transformed fluorescence of one flow cytometry data file. We excluded the fluorescence distributions of start codon mutations for reasons discussed in the main text.

**Table S3:** Parameters for the computational model of fitness obtained from flow cytometry data for differentially enriched mutations using **(A)** green and **(B)** cyan fluorescence measurements.

| Parameter | Mean | Std. deviation |
| --- | --- | --- |
| $\mu'_{+L}$ | 0.9351 | 0.0234 |
| $\mu'_{+H}$ | 1.0729 | 0.0294 |
| $C_{+L}$ | 0.3961 | 0.0297 |
| $C_{+H}$ | 0.5941 | 0.0321 |
| $\sigma_{+L}$ | 0.1623 | 0.0129 |
| $\sigma_{+H}$ | 0.1702 | 0.0074 |
| $\sigma_{-}$ | 0.1860 | 0.0199 |

| Parameter | Mean | Std. deviation |
| --- | --- | --- |
| $\mu'_{+L}$ | 0.9312 | 0.0273 |
| $\mu'_{+H}$ | 1.0782 | 0.0286 |
| $C_{+L}$ | 0.3954 | 0.0304 |
| $C_{+H}$ | 0.6002 | 0.0301 |
| $\sigma_{+L}$ | 0.1604 | 0.0118 |
| $\sigma_{+H}$ | 0.1717 | 0.0074 |
| $\sigma_{-}$ | 0.1856 | 0.0207 |

**Figure S17:**
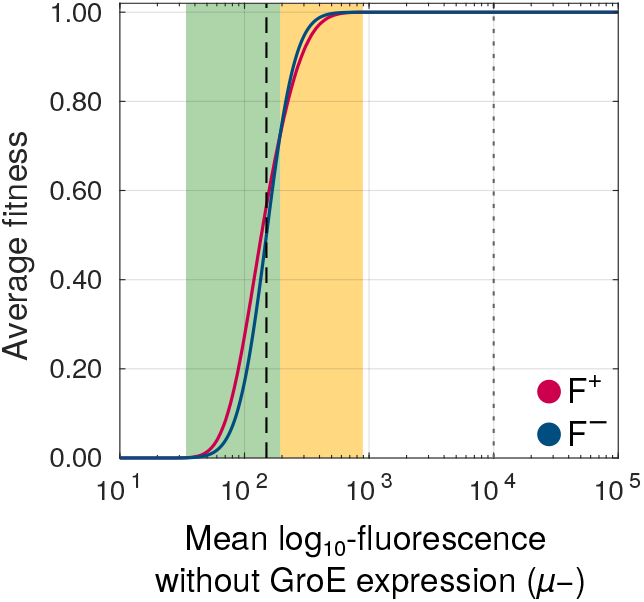
The average fitness (vertical axis) of variants with (red) and without (blue) GroE expression increases with increasing values of μ- (horizontal axis). The selection threshold we used during experimental evolution (150 arbitrary units of fluorescence) is denoted by the dashed black vertical line. Despite a steady increase in fitness with increasing fluorescence under both conditions, GroE increases fitness of some genotypes (green area), reduces the fitness of some other genotypes (orange area), while not affecting the rest (white area). Ancestral GFP is denoted by the grey dotted vertical line. See also Main text: Figure 6.

## 12 List of primers

**Table S4:**
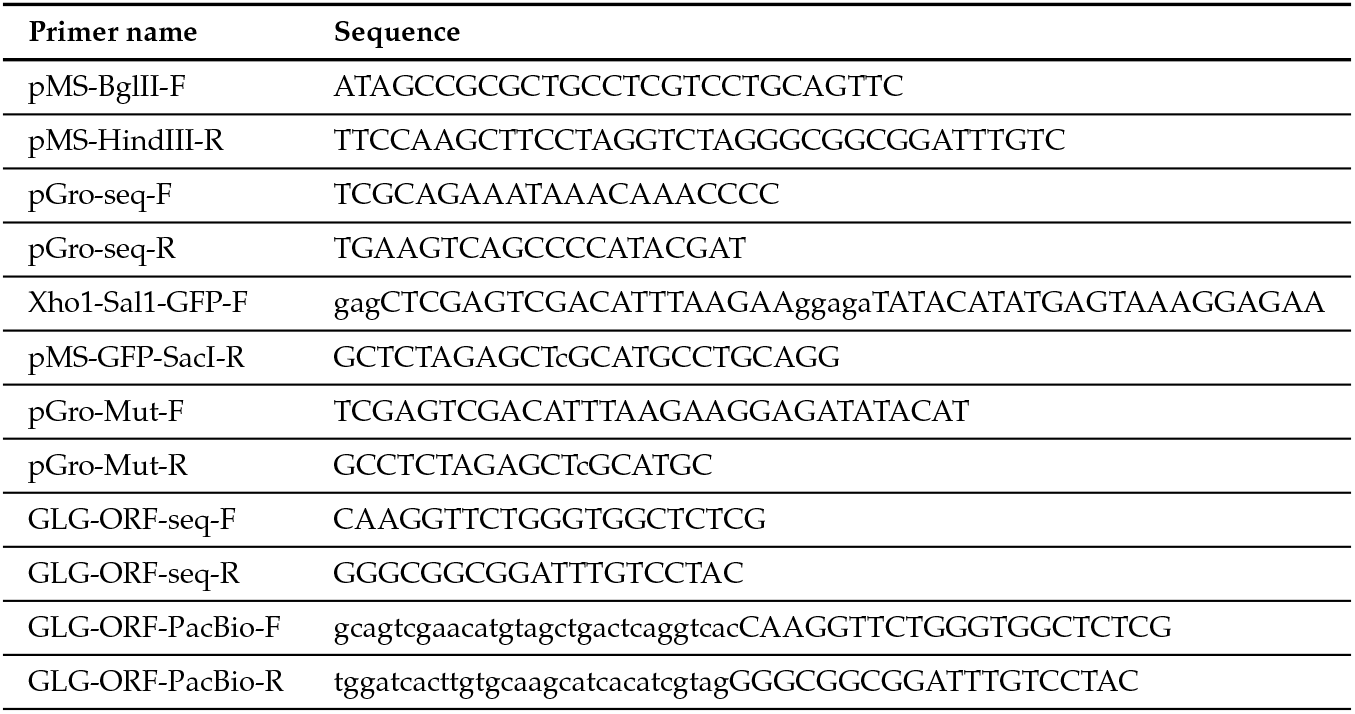
List of primers used for cloning, PCR based random mutagenesis and SMRT sequencing library preparation.

**Table S5:**
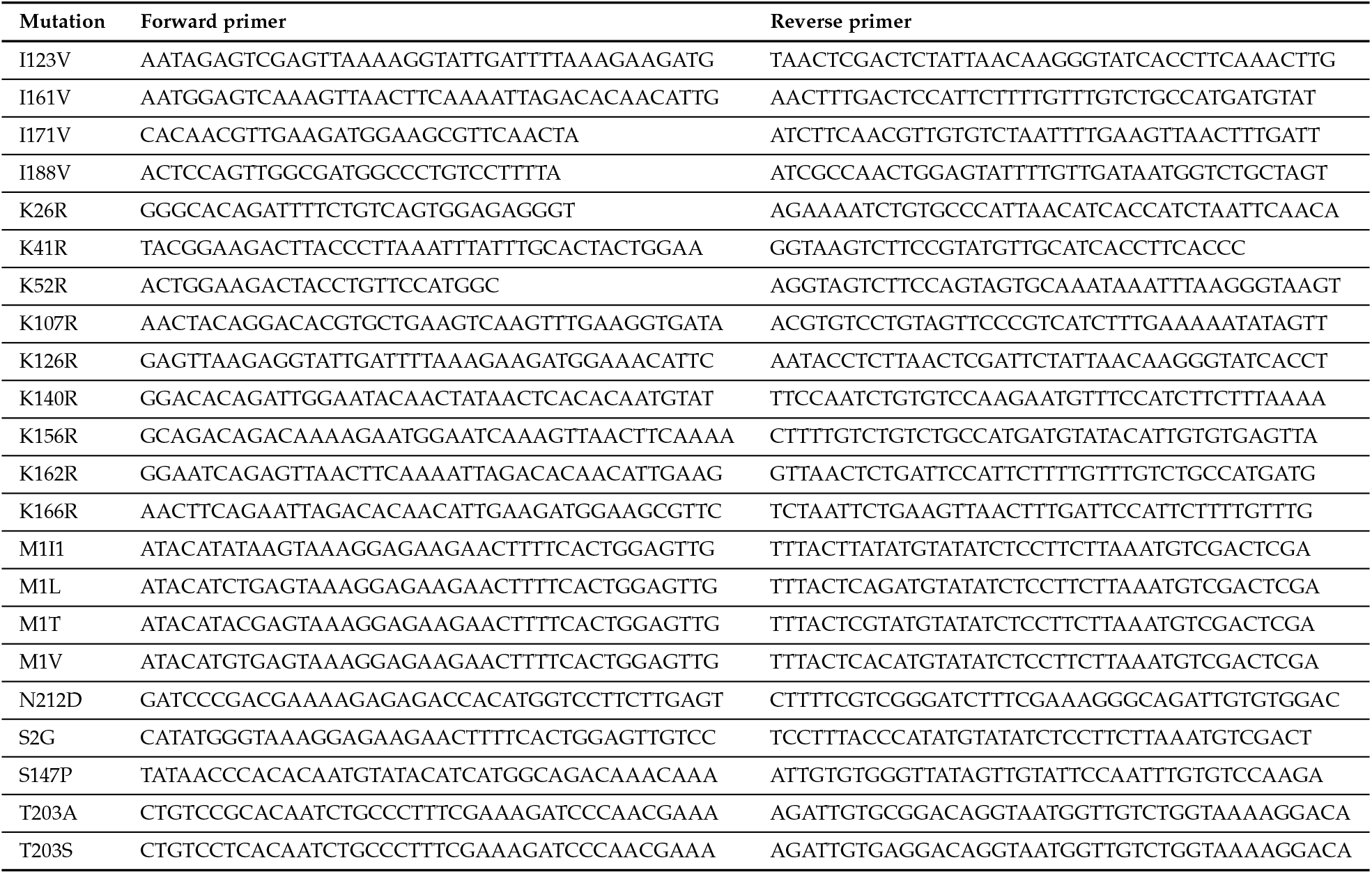
List of site directed mutagenesis primers.

